# Subcellular mRNA kinetic modeling reveals nuclear retention as rate-limiting

**DOI:** 10.1101/2024.03.11.584215

**Authors:** David Steinbrecht, Igor Minia, Miha Milek, Johannes Meisig, Nils Blüthgen, Markus Landthaler

**Author notes:** joint first authors (David Steinbrecht, Igor Minia).

## Abstract

Eukaryotic mRNAs are transcribed, processed, translated, and degraded in different subcellular compartments. Here, we measured mRNA flow rates between subcellular compartments in mouse embryonic stem cells. By combining metabolic RNA labeling, biochemical fractionation, mRNA sequencing, and mathematical modeling, we determined the half-lives of nuclear pre-, nuclear mature, cytosolic, and membrane-associated mRNAs from over 9000 genes. In addition, we estimated transcript elongation rates. Many matured mRNAs have long nuclear half-lives, indicating nuclear retention as the rate-limiting step in the flow of mRNAs. In contrast, mRNA transcripts coding for transcription factors show fast kinetic rates, and in particular short nuclear half-lives. Differentially localized mRNAs have distinct rate constant combinations, implying modular regulation. Membrane stability is high for membrane-localized mRNA and cytosolic stability is high for cytosol-localized mRNA. mRNAs encoding target signals for membranes have low cytosolic and high membrane half-lives with minor differences between signals. Transcripts of nuclear-encoded mitochondrial proteins have long nuclear retention and cytoplasmic kinetics that do not reflect co-translational targeting. Our data and analyses provide a useful resource to study spatiotemporal gene expression regulation.

## Introduction

The life cycle of mRNA is a complex process involving multiple steps in different subcellular compartments (Glisovic *et al*, 2008). The precise regulation of mRNA dynamics in each step is critical for cellular transcript homeostasis (Berry & Pelkmans, 2022). In a typical mammalian cell, about 10.000 protein-coding genes are expressed, producing thousands of mRNAs per minute (Schwanhäusser *et al*, 2011). Protein-coding transcripts undergo maturation in the nucleus through splicing and polyadenylation, followed by export to the cytosol. Translation either occurs in the cytosol or at membrane-bound organelles, predominantly the endoplasmic reticulum (ER), culminating in their turnover. The resulting flow of mRNAs affects cell function by shaping the dynamic amount of mRNAs available for translation in the cytoplasm (Eisen *et al*, 2020; Mor *et al*, 2010) and the ER (Das *et al*, 2021). The overall mRNA abundance scales with cell size (Padovan-Merhar *et al*, 2015; Kempe *et al*, 2015; Battich *et al*, 2015; Swaffer *et al*, 2023).

Notably, mRNA half-lives exhibit a remarkable variability exceeding 100-fold among different protein-coding transcripts (Herzog *et al*, 2017; Dölken *et al*, 2008; Friedel *et al*, 2009). To unravel the nuanced regulation of individual processes in the mRNA life cycle, a crucial need arises for time-resolved and subcellular quantification.

Early studies on the subcellular dynamics of mRNA considered either all poly(A) RNA collectively (Jelinek *et al*, 1973) or only a few individual genes (Mor *et al*, 2010; Grünwald *et al*, 2011; Tutucci *et al*, 2018). Recent technological advances have enabled global measurements of mRNA dynamics and subcellular distribution. Metabolic labeling of newly synthesized RNA with uridine analogs has provided experimental means to monitor RNA dynamics with a minimal perturbation to the cell (Dölken *et al*, 2008). Using 4-thiouridine for labeling, kinetic rates of mRNA have been quantified on a transcriptome-wide level in a number of cell types from different species (Chen & van Steensel, 2017; Herzog *et al*, 2017; Rabani *et al*, 2011; Rutkowski & Dölken, 2017; Schofield *et al*, 2018). The first global measurement of global nucleocytoplasmic mRNA dynamics was achieved by Chen and colleagues in Drosophila S2 cells (Chen & van Steensel, 2017). However, metabolic labeling studies on mammalian cells have mostly considered the total cellular mRNA to determine kinetic rates, overlooking subcellular aspects (Herzog *et al*, 2017; Rabani *et al*, 2011; Rutkowski & Dölken, 2017). A recent study based on single-cell in-situ sequencing of 5-ethynyl uridine-labeled RNA measured transcription, translocation and degradation of individual transcript molecules, uncovering subcellular mRNA profiles across time and space at the single-cell level for a collection of almost 1000 genes (Ren *et al*, 2023). Furthermore, two independent works combined 4-thiouridine, cellular fractionation and RNA sequencing to measure the rates at which RNAs are exported from the nucleus in mammalian cells (preprint: Müller *et al*, 2023; preprint: Smalec *et al*, 2022), with one study additionally providing evidence for mRNA degradation in the nucleus (preprint: Müller *et al*, 2023).

Here, we combined metabolic RNA labeling with cellular fractionation of mouse embryonic stem cells in nuclear and cytosolic, also a membrane-bound fraction composed mostly of the ER and mitochondria to produce mRNA sequencing data with spatial and temporal resolution. We developed a mathematical framework that enabled us to infer the kinetic rate constants of nuclear pre-, nuclear mature, cytosolic and membrane-bound mRNAs transcriptome-wide from our metabolic labeling and fractionation time series sequencing data. In addition, we estimated a transcript elongation rate to account for an observed labeling incorporation bias. Our method provides subcellular half-life information of the relative contributions of each step of the mRNA life cycle to transcript steady-state levels. We provide evidence that nuclear retention of mRNAs is the rate-limiting step in the life cycle of protein coding transcripts. In addition, it uncovers expected differences in rates of mRNAs coding for distinct protein families, such as transcription factors and proteins of the secretory pathway. Comparison of the kinetic rate constants for these steps across genes will provide novel insights into subcellular mechanisms of differential gene regulation.

## Results

### Spatiotemporal measurement of newly-transcribed mRNA

To capture the nucleocytoplasmic kinetics of mRNA, we conducted a time-resolved SLAM-seq experiment in combination with subcellular fractionation in mouse embryonic stem cells (mESCs). Briefly, mESCs were exposed to 4-thiouridine (4sU) for various times to label newly synthesized RNA. Subsequently, we performed subcellular biochemical fractionation to generate cytosolic, membrane and nuclear fractions (Fig. 1A). 4sU concentrations were carefully chosen for sufficient labeling rates while minimizing a potential toxicity elicited by 4sU, with higher concentrations of 4sU (500 µM) for labeling periods from 15 minutes to 3 hours, and lower concentrations (100 µM) from 1 to 3 hours. A differential gene expression analysis confirmed that 4sU labeling caused none or minimal perturbation of transcriptome (Sup. Fig. 1A). The efficacy of the fractionation procedure was confirmed by Western blot analysis with antibodies against compartment-specific marker proteins. The nuclear proteins histone H3, lamin A/C (LMNA) and TATA-binding protein (TBP), the cytoplasmic proteins GAPDH and β-tubulin (TUBB) and the ER marker protein BCAP31 were only detectable or strongly enriched in their respective fraction. In contrast, RPS6, a component of the 40S ribosomal subunit, was found in all fractions (Fig. 1B).

**Figure 1:**
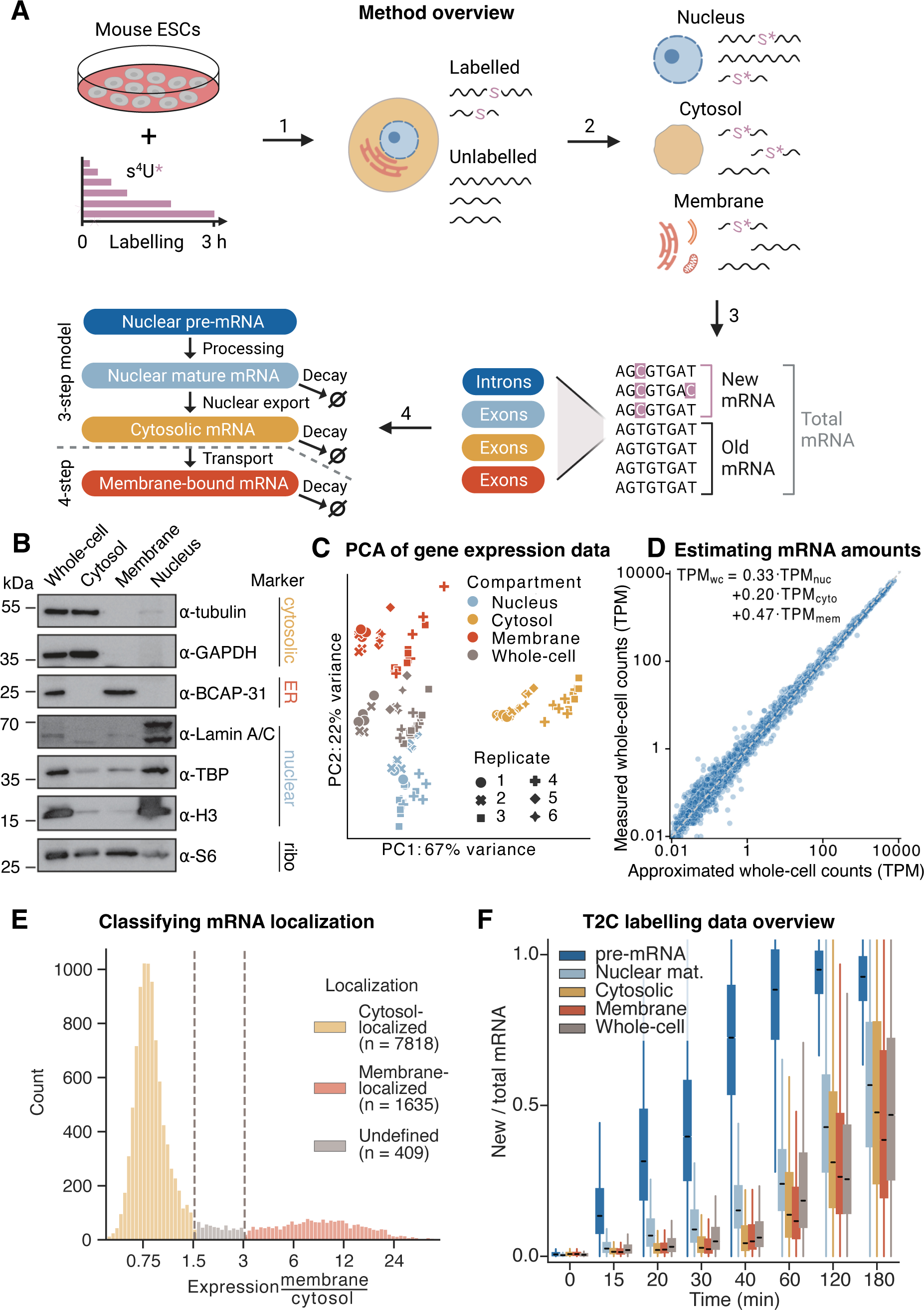
Newly-transcribed mRNA is measured spatio-temporally using cell fractionation and metabolic labeling RNA sequencing. **(A)** Schematic overview of experimental and computational steps. Mouse embryonic stem cells (mESCs) are labeled with 4sU for 15, 20, 30, 40, 60, 120 and 180 minutes, leading to T2C conversions in newly-transcribed RNA (1). Cells are fractionated into nuclear, cytosolic and membrane-bound compartments (2). Ratios between new and total mRNA are estimated in a Bayesian framework for exons and whole intronic regions (3) and given as input to a kinetic model to fit subcellular kinetic rate parameters (4). See Methods for details on each individual step. **(B)** Results from Western blot analysis measuring nuclear, cytosolic, endoplasmic reticulum and ribosomal protein markers in the subcellular fractions and the whole cell extract. (**C)** Principal component analysis of log-transformed bulk mRNA-seq data in transcripts per million (TPM) using 2,000 most variable genes. Samples cluster by compartment. **(D)** Estimates of relative mRNA abundance in subcellular compartments. Subcellular TPM values are fitted to whole-cell TPM values, resulting in estimates for the relative abundance of mRNA in each compartment. **(E)** Classification of membrane and cytosolic mRNA localization in mESCs through subcellular mRNA expression. Histogram of the ratio between steady-state gene expression (TPM, mean over all time points) in membrane and cytosolic fractions, from here on membrane enrichment, for 11,500 most expressed genes. Vertical gray lines indicate chosen cutoffs to classify mRNA localization. For membrane enrichment < 1.5, < 3 and > 3: cytosol-localized, undefined and membrane-localized, respectively Number of successfully fitted genes in each localization category is shown in the legend. **(F)** Time- and compartment-resolved box plot of T2C labeling data. Medians of share of new to total mRNA increases with labeling time. Lower and upper hinges of box plots correspond to the 25th and 75th percentiles, respectively. Lower and upper whiskers extend from the hinge to the smallest or largest value no further than the 1.5x interquartile range from the hinge, respectively. Center lines of box plots depict the median values.

To quantify the subcellular kinetics of mRNA globally, we next extracted RNA from each fraction and from whole cells, followed by iodoacetamide (IAA) alkylation and poly(A)-selected strand-specific mRNA library preparation. Briefly, sequencing reads were processed, aligned to the mouse genome and quantified for gene expression, T and T2C conversion counts in both exons and introns (see Methods for details). Based on a binomial mixture model (Jürges *et al*, 2018) we determined the conversion rate per sequenced RNA sample. With the conversion rates, T and T2C conversion counts the share of new to total mRNA was calculated on intron and exon level, which was used later on as an input for the kinetic model (see Fig. 1A bottom).

The experimental and computational complexity demanded an evaluation of the effects of biological and technical variability on mRNA quantification. Principal component analysis (PCA) revealed that the largest differences in mRNA quantification were a consequence of the assayed subcellular compartment rather than the difference in biological replicates (Fig.1C), indicating that the data captured biological variability well with a low influence of technical effects.

To convert relative to absolute subcellular RNA expression ratios, we needed to quantify how the total amount of RNA is shared between the subcellular compartments. Under the assumption that the RNA expression in the whole cell can be reconstructed by summing the subcellular RNA expression with a corresponding factor for each, we fit the relative abundance of nuclear, cytosolic and membrane mRNA with the constraint that they sum up to 1 (the whole-cell expression). We estimated that 33%, 20% and 47% of total mRNA is present in the nucleus, cytosol and membrane-bound, respectively (see Fig. 1D). These relative abundances are used to derive expression ratios mimicking absolute RNA levels that are related to kinetic parameter ratios and can therefore be used to constrain the parameter space (see Box 1).

Comparing gene expression in the cytosol and the membrane, we observed a bimodal distribution, with 16% and 79% of the transcripts being enriched at the membrane and in the cytosol, respectively (Fig. 1E). Based on this distribution, we classified mRNAs into cytosol- and membrane-localized and a small group of undefined transcripts not belonging to these compartments. This classification determined the particular model used in the fitting process (see bottom left Fig. 1A).

In addition, the distribution of share of labeled mRNAs over all quantified genes (n=9862) reflected the cellular mRNA maturation and showed expected trends for all subcellular compartments (Fig.1E). Collectively, these results showed that the quantification of global mRNA kinetics in subcellular fractions of mouse ES cells was of high quality.

### Model of subcellular mRNA dynamics

To integrate the subcellular transcriptome and metabolic RNA labeling data and to estimate half-lives as a description of subcellular mRNA dynamics, we developed transcript-wise mathematical models similar to previous work (28771467). Each transcript was modeled by a linear, inhomogeneous system of ordinary differential equations (ODE) containing three or four steps for cytosol- or membrane-localized transcripts, respectively, describing the life cycle from mRNA transcription to degradation (see Box 1). After transcription in the nucleus, nuclear pre-mRNA is processed to mature mRNA (pre-mRNA processing rate *k*_1_). From there, mature mRNA can be exported to the cytosol or already be degraded (nuclear export *k*_2_ and nuclear degradation *γ*_2_). In the 3-step model, mRNA is translated and degraded in the cytosol (cytosolic decay *γ*_3_), whereas in the 4-step model, mRNA is first localized to the ER membrane or other membrane-bound organelles (cytosolic transport *k*_3_, no decay), where it is translated and subsequently degraded (membrane decay *g*_4_). We derived a system of ODEs describing the relative amount of new mRNA. The share of new to total mRNA estimated from the SLAM-seq data is used as input in the solutions of this ODE system to fit the parameters mentioned above.

For ease of interpretation, we will mainly use half-lives instead of rates in both text and figures in the following (see Box 1), namely: pre-mRNA processing (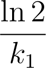), nuclear retention (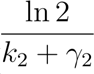), cytosolic stability (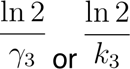) and membrane stability (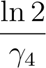). Nuclear retention time is used as a general term to comprise the various events after maturation of a single transcript molecule until its emergence into the cytoplasm, including chromatin dissociation, nuclear diffusion, and binding to and transport across the nuclear pore. Figures that include both the elongation rate and subcellular kinetic parameters display the corresponding rates (in kb · min^-1^ and min^-1^, respectively).

#### Box 1.

##### Kinetic model of the mRNA life cycle

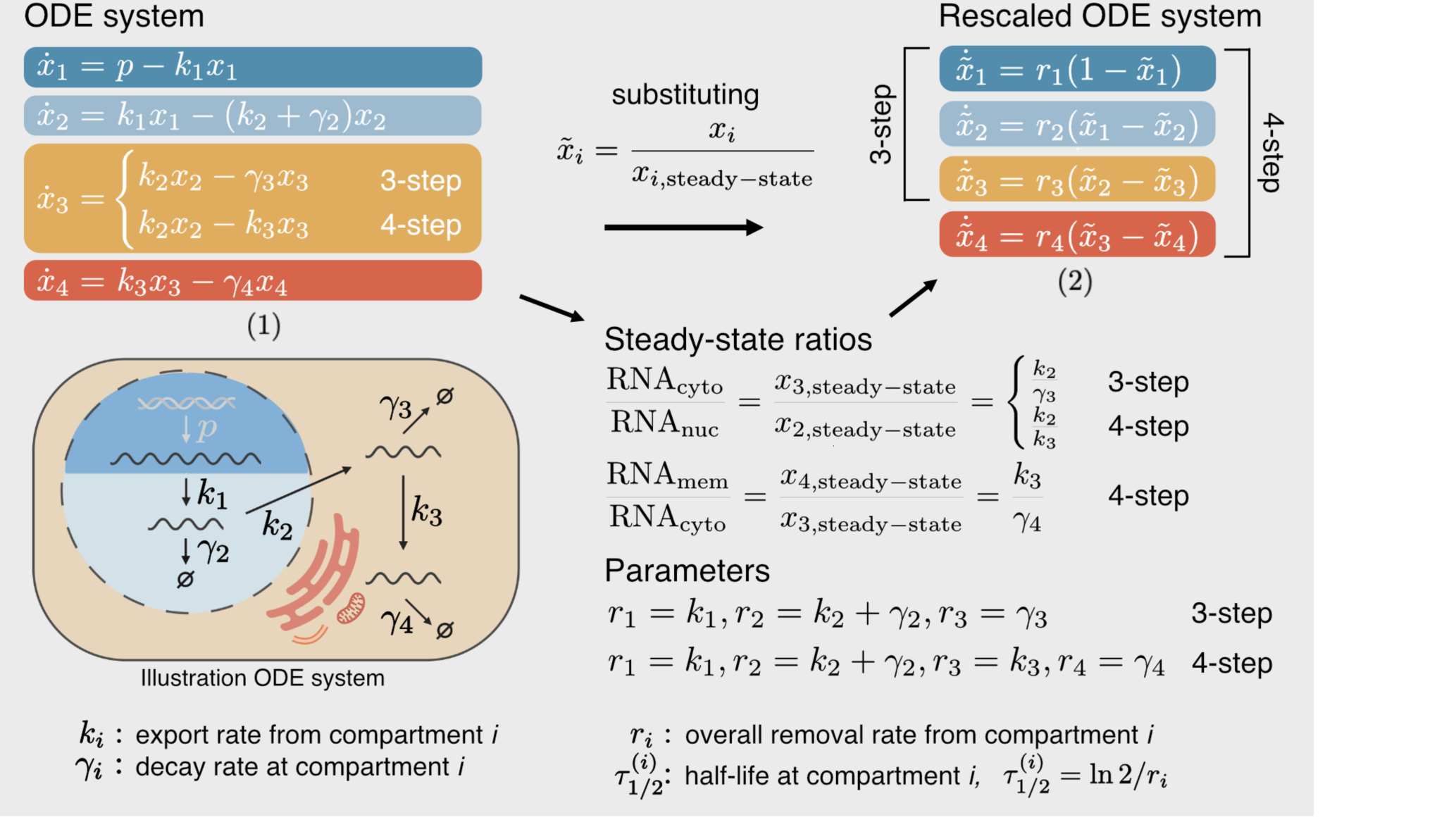

The individual equations in Eqs. (1) describe the change in absolute amount of new nuclear pre- (dark blue), nuclear mature (light blue), cytosolic (yellow) and membrane-bound (red) mRNA. The biological process of each parameter is illustrated in the sketch on the bottom left. We rescale the system by dividing each compartment with its steadystate solution, so that Eqs. (2) describe the change in relative share of new mRNA. The analytical solutions to Eqs. (2) are shown in Box 2 in the methods. We use the solutions of the first three (3-step model) and all four equations (4-step model) to fit cytosol- and membrane-localized mRNAs, respectively. The steady-state solutions additionally constrain the parameter space, allowing to distinguish nuclear export from decay using the measured subcellular RNA expression (see steadystate ratio equations). The production rate of pre-mRNA p is not present in the rescaled system and hence not fitted.

### Modeling of transcriptional elongation rates

For mRNAs whose transcription is initiated prior to labeling with 4sU, but terminated after labeling start, we expected that only part of the transcript was labeled. More specifically, we expect that sequences at a distance of d to the 3’ end are only labeled after *t* = *d/v*, where *v* is the transcription elongation rate (see Fig. 2A). Therefore, particularly for longer transcripts and shorter time points, partial labeling affects our measurements and has to be taken into consideration.

**Figure 2:**
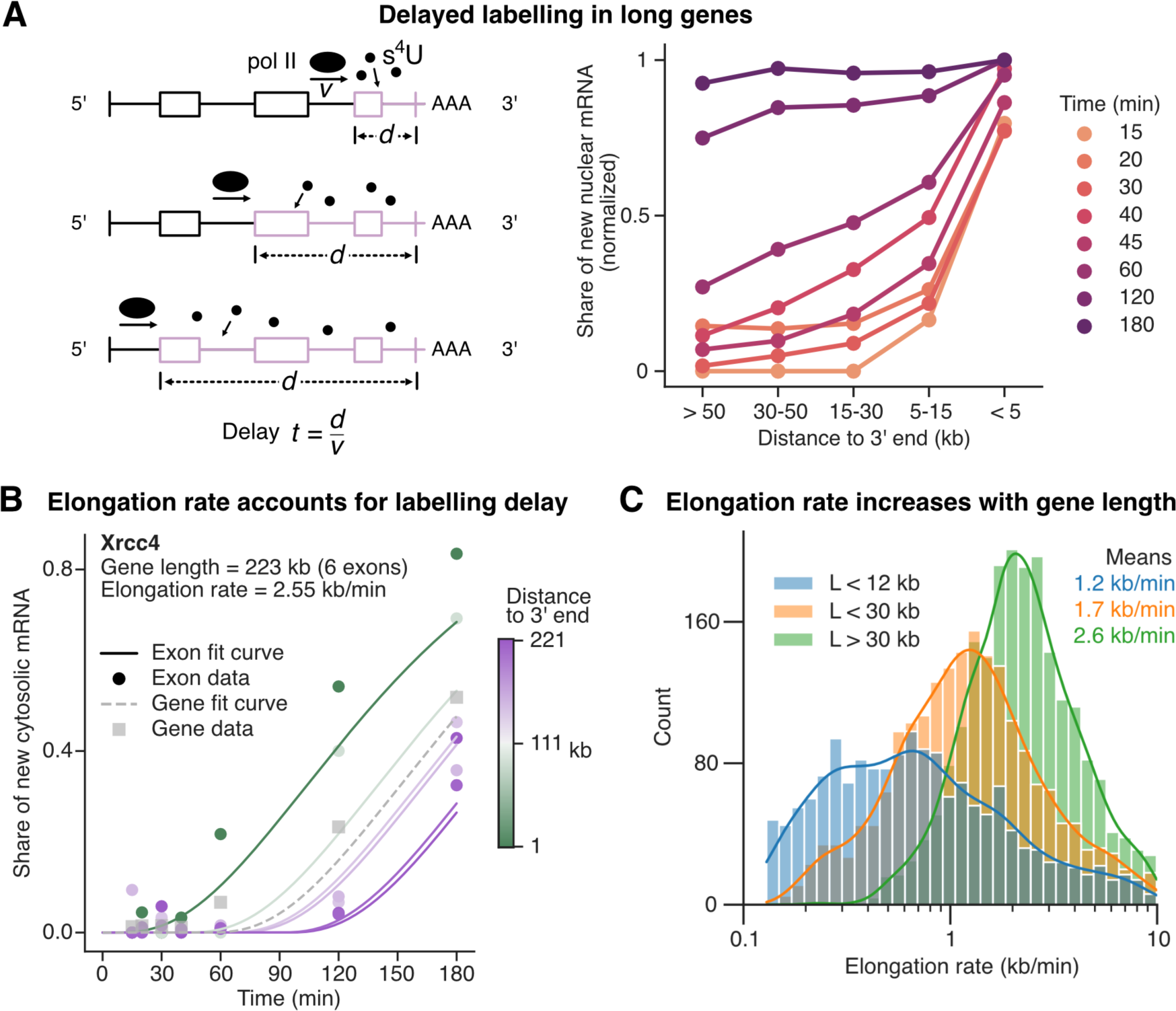
Transcript elongation rate is estimated to account for labeling delay in long genes. **(A)** Left: Sketch to illustrate how exons (in longer genes) are labeled with a time-delay that increases linearly with distance to 3’ end. As poly(A)+ sequencing was used, transcripts close to being fully transcribed will contain T2C conversions first (close to the 3’ end), while it takes longer to observe conversions in the whole transcript. Right: Pointplot of new over total exonic, nuclear mRNA binned by distance to 3’ end. Points show medians across exons per bin and time and are normalized to the 15 min and < 5 kb point. Lines connect points of equal time. Equal labeling efficiency is seen after 3 hours. **(B)** Fit result of *Xrcc4* (only cytosolic mRNA) is shown to illustrate the fitting procedure. All exons of a gene share the same kinetic parameters, but each exonic curve is time-delayed by *t̃* = *t* – *d/v* – *δ*, with *d* being the mean exonic distance to 3’ end, *v* the elongation rate and the overall delay of 5 min (straight lines). The closer an exon is to the 3’ end (indicated by color), the higher its ratio of new mRNA is (points). The time-delayed fit accounts for that. Gray squares show the gene-level T2C data. Gray, dashed line shows a fit curve delayed by mean 3’ end distance weighted by expression of exons. **(C)** Histogram with kernel-density estimation of reliably estimated elongation rates (n = 5661) with genes grouped and colored by gene length (L). Mean rate per group is shown in the top right. Elongation rate tends to increase with gene length.

Our transcriptome sequencing data covers poly-adenylated RNA, and quantification of labeling includes sequences spanning the entire transcripts. When we grouped exonic sequences by distance to the 3’ end, we observed that the median share of labeled mRNAs in the nuclear fraction decreased with increasing distance to the 3’ end, particularly for shorter time periods (see Fig. 2A). To account for this bias, we incorporate a transcript elongation rate in our model. Specifically, we modeled each transcript with one variable per exon, where the dynamics of each exon is modeled using the model described above but with a time-delay according to *t̃* =.*t* – *d/v* with d being the distance of the exon to the 3’ end (see Fig. 2B). All exon models of one transcript share the same kinetic parameters.

### Model parameterization unveils length-dependent elongation rate

To derive estimates of kinetic parameters and elongation rates, we fitted the models to the time-resolved SLAM-seq data. More specifically, we simultaneously optimized model parameters for each transcript such that they best fit (a) the share of labeled mRNA at the level of each exon for the different time points and (b) the steady-state levels of the transcripts in the nucleus, cytoplasm and membrane compartment. Out of 8548 transcribed genes with sufficient labeling data in multiple exons, we could reliably estimate the elongation rate for 5661 genes with a mean of 1.9 kb · min^-1^. For gene lengths shorter than 12 kb, between 12 kb and 30 kb and longer than 30 kb, the mean elongation rate is 1.2, 1.7 and 2.6 kb · min^-1^, respectively (see Fig. 2C). Increases in elongation rate of RNA polymerase II for longer genes, as well as our quantitative range of values, coincide with results from previous studies focusing specifically on transcript elongation (Jonkers *et al*, 2014; Fuchs *et al*, 2014; Veloso *et al*, 2014).

### Nuclear retention time is the rate limiting step in the life cycle of most mRNAs

For mRNAs derived from roughly 9,800 genes, we observed three common kinetic profiles: i) for transcripts with fast turnover labeled RNA dynamics are similar in all compartments (exemplified by *Myc*, Fig. 3A), with the exception of nuclear pre-mRNA; ii) cytosol-stable transcripts show a slower accumulation of labeled RNA in the cytosolic than in the nuclear mature compartment (e.g. *Nf1*, Fig3A), with overall turnover varying from fast to slow; iii) membrane-localized mRNAs, as defined earlier, accumulate labeled RNA equally fast in nuclear mature and cytosolic compartments, but significantly more slowly in the membrane than in the cytosolic compartment (e.g. *Tfrc*, Fig.3A). This distinct pattern is due to the low cytosolic expression and the short cytosolic residence time of these transcripts.

**Figure 3:**
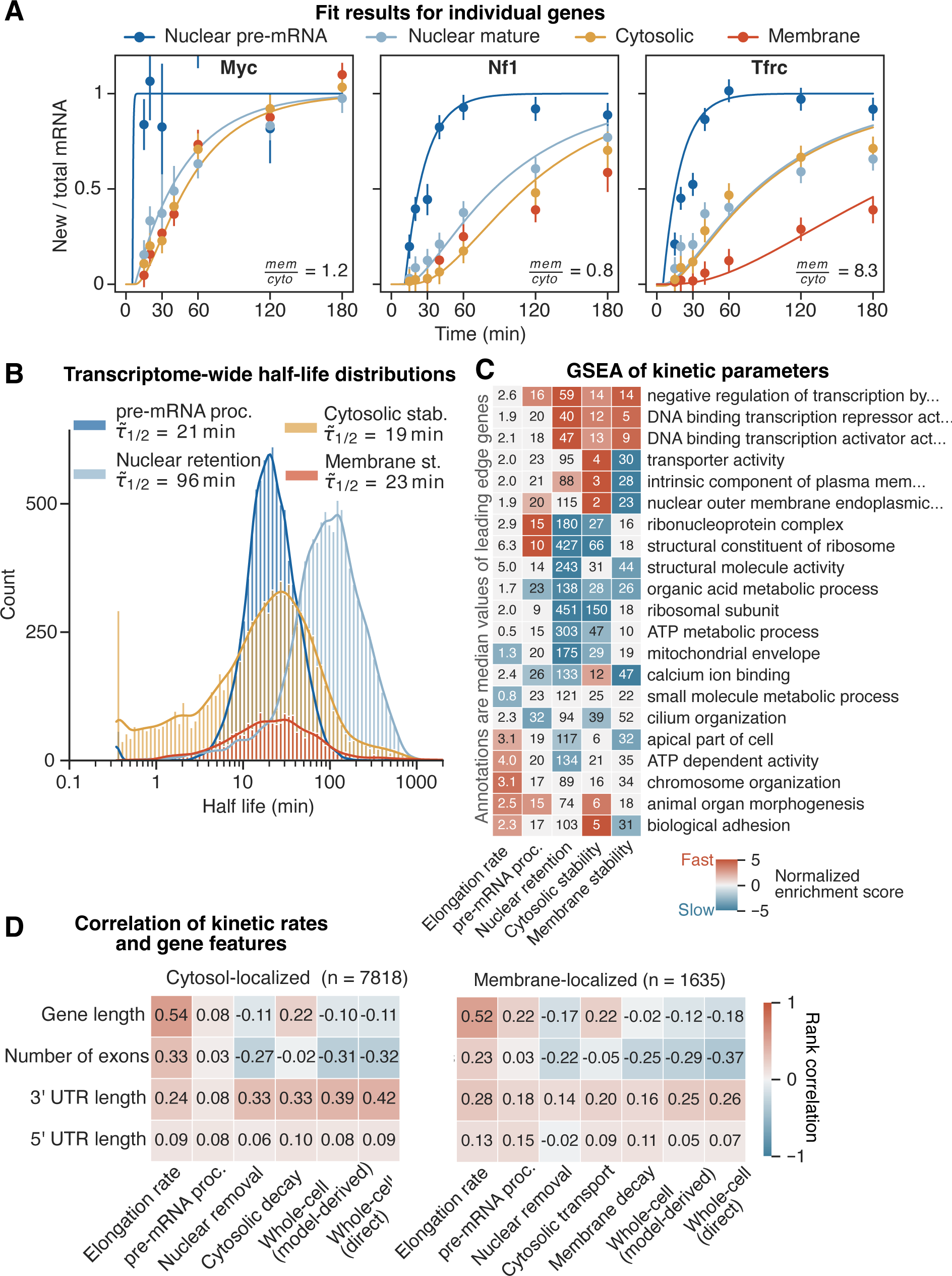
Across the transcriptome, transcripts show a wide distribution and different combinations of subcellular kinetic parameters. **(A)** Fit results of three exemplary genes. Ratio of new to total mRNA is shown over time. Points with error bars are mean with standard deviation across replicates per fraction and time (all fractions shown). Lines are fit results, delayed as the gene-level curve in Fig. 2B (only fitted compartments shown). Both points and lines are colored by compartment. Ratio between steady-state expression in membrane and cytosolic compartments is shown in bottom right, indicating if 3-step or 4-step model was used (< 1.5 or > 1.5, respectively, see Fig. 1E). **(B)** Histogram of nuclear pre-mRNA processing (n = 8548), nuclear retention (n = 9862), cytosolic stability (n = 9862) and membrane stability (n = 2044) mRNA half-lives. Bars are colored by parameter. Median half-life of each parameter is shown in legend. **(C)** Heatmap of results from five GSEAs based on the gene ontology (GO). Genes were ranked by log-scaled, z-scored parameter rates. Rows are GO terms, columns are kinetic parameters. For each parameter, the two most up- and down-regulated terms are shown. Additionally, five terms with the lowest adjusted p-values and adjusted p-values < 0.05 in at least three columns are shown. Color bar depicts the value of a normalized enrichment score. Gray indicates the term is not significant. Values in heatmap are median half-lives (rate in the first column) of the union of leading edge genes across each row. **(D)** Heatmaps of correlation between kinetic rates and gene features. Model-derived whole-cell half-lives are calculated from subcellular rates, see Supp. Fig. 4A and Methods. Direct whole-cell half-lives obtained by fitting a simple “one minus exponential decay” model to the share of new to total mRNA from whole-cell extract data. Transcripts are split into cytosol-(left) and membrane-localized (right). Values in heatmap are spearman rank correlations, also indicated by color bar.

All kinetic parameters for transcripts in different compartments were highly variable (see Fig. 3B). Nuclear pre-mRNA half-life showed the narrowest distribution, ranging from 9 to 48 min (10th and 90th percentile, resp.) with a median of 21 min. Cytosolic transcript half-lives displayed the widest distribution and the highest variability, ranging from 1 to 84 min (10th and 90th percentile, resp.) with a median of 19 min. Membrane mRNA half-lives range from 5 to 91 min with a median of 23 min. Interestingly, our data suggest that the nuclear retention of mature mRNAs is the rate limiting step for most transcripts with a median half-life of 96 min and ranging from 28 min to 290 min. This observation agrees with findings by Mueller and colleagues describing the nucleus-to-cytosol step to be rate-limiting (preprint: Müller *et al*, 2023).

In summary our analysis suggests that most mRNAs spend more than half of their lifetime in the nucleus as mature spliced transcripts. Recent results using metabolic labeling RNA sequencing suggested that nuclear transcripts remain associated to DNA for a longer time, and once dissociated they are exported rather quickly (preprint: Smalec *et al*, 2022).

### Functionally related transcripts tend to have similar kinetic profiles

It has been previously shown that groups of mRNAs encoding proteins with specific molecular functions display distinct kinetics rates along the mRNA life cycle (Chen & van Steensel, 2017; Ren *et al*, 2023). To this end, we performed gene set enrichment analyses (GSEA) considering the pre-mRNA processing, nuclear retention, cytosolic and membrane stability, respectively, to identify groups of transcripts from the gene ontology (GO) that have particularly short or long residence time in each compartment. Each GSEA resulted in a large number of significantly enriched GO terms, with a subset shown in Fig. 3C. For the nuclear retention time, mRNAs encoding transcription regulators have a high positive enrichment score, i.e. short nuclear retention time, while those involved in metabolic processes, specifically translation, have a high negative enrichment score, i.e. a long nuclear retention time. Most transcripts of mitochondrial and ribosomal genes show particularly long nuclear half-lives. This may mechanistically be explained by their enrichment in nuclear speckles, where mRNA transcripts of both gene groups were found to be retained and post-transcriptionally processed (preprint: McIntyre *et al*, 2023). Transcripts associated with transport or ion homeostasis show short cytosolic and long membrane half-lives. When focussing on membrane-localized transcripts, those associated with regulating protein phosphorylation showed the shortest (∼9 min), while those associated with calcium ion binding showed the longest (∼61 min) membrane half-lives (see Supp. Fig. 3C). Next, we investigated if kinetic parameters were correlated with gene features including gene length, number of exons, and length of UTRs. The strongest association was observed with the transcriptional elongation rate, as noted above, and presented a rank correlation of more than 0.5 with gene length (see. Fig. 3D). Nuclear retention and whole-cell half-lives are both positively correlated with the number of exons in both cytosol- and membrane-localized transcripts (see Fig. 3D). Therefore, mRNAs with many exons take longer to be exported but are then overall more stable. However, for cytosol-localized transcripts, the cytosolic half-life is not correlated with exon number, while the membrane half-life for membrane-localized genes is. Interestingly, the number of exons seems to play an important role in subcellular mRNA kinetics. mRNA derived from longer genes seem to be degraded faster in the cytosol, but not at the ER (Fig. 3D). 3’ UTR length correlates moderately for cytosol- and weakly for membrane-localized mRNAs with all subcellular parameters except pre-mRNA processing, suggesting that the longer the 3’ UTR end, the shorter the half-life as described previously (Spies *et al*, 2013). Out of the subcellular parameters, nuclear retention is the best predictor of whole-cell half-life (Spearman correlation of r = 0.88 and r = 0.8 for model-derived and directly-fitted whole cell half-life, respectively).

### Differentially localized mRNAs exhibit distinct dynamic behavior

To associate mRNA localization with function, we ranked transcripts by membrane enrichment, which was also used to classify mRNA localization, and performed a GSEA. The proteins encoded by transcripts highly enriched in the membrane fraction are located at the plasma membrane, cell surface and endoplasmic reticulum and are involved in transmembrane transport and ion homeostasis (Fig. 4A). Proteins of the more than 7000 cytosol-localized transcripts are distributed across the cell and have a wide range of biological functions, but those encoded by transcripts most highly enriched in the cytosol are located primarily in the nucleus and are involved in organization of chromatin and chromosomes and transcription regulation (Fig. 4A).

**Figure 4:**
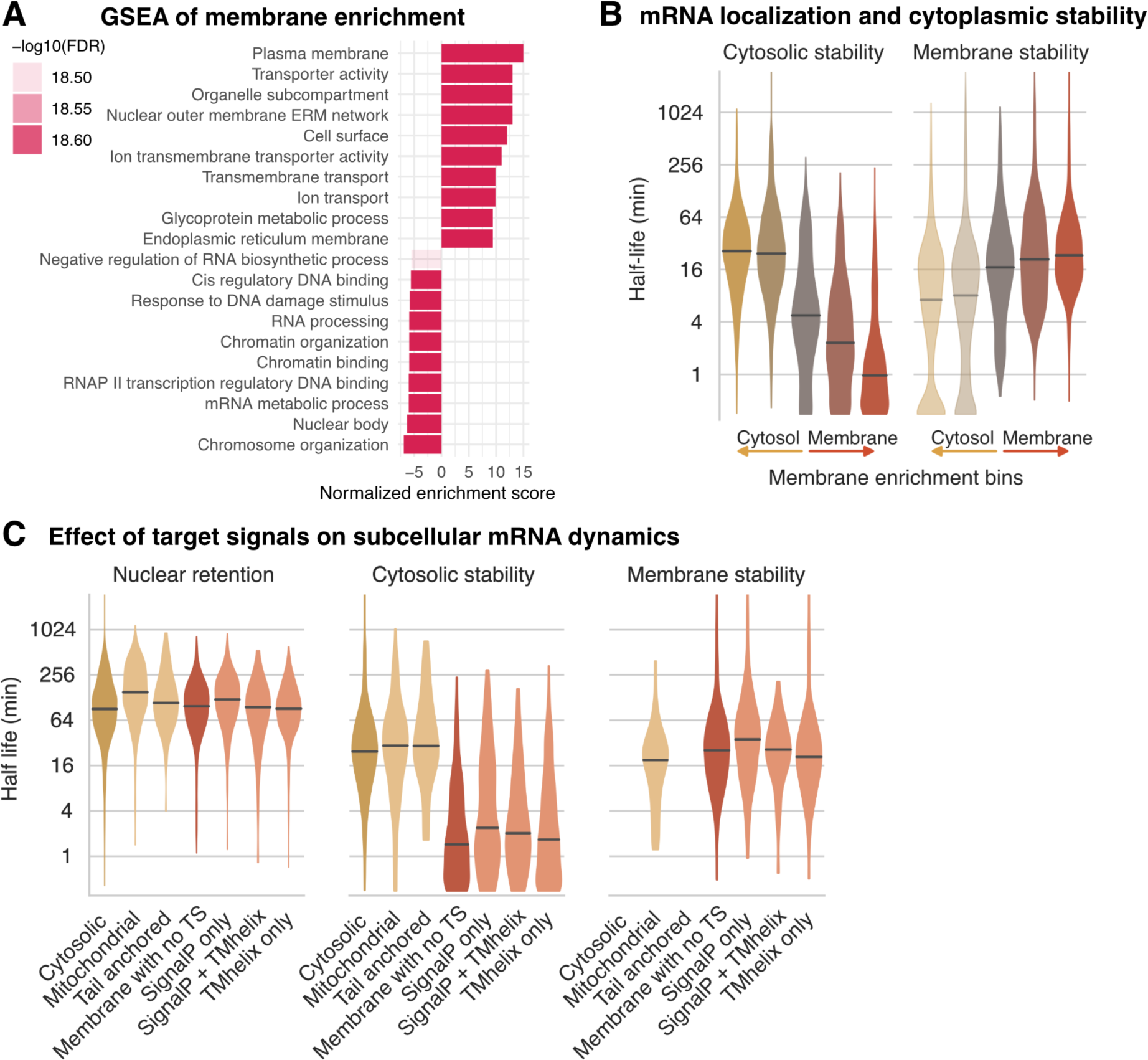
Differentially localized mRNAs exhibit distinct combinations of kinetic rate constants. **(A)** Result of a GSEA ranking genes by the ratio of membrane over cytosolic RNA expression (membrane enrichment). Color transparency shows Benjamini-Hochberg corrected falsediscovery rates (FDR). **(B)** Violin plot of cytosolic and membrane half-lives with transcripts binned by membrane enrichment (see Fig. 1E with cytosol- and membrane-localized further split at 0.8 and 8.7, respectively). Yellow, gray and red colors indicate cytosol, undefined and membrane localization, respectively. Membrane half-lives of cytosol-localized transcripts from additional 4-step model fit are shown transparently. Center lines of violin plots depict the median values. **(C)** Violin plot of nuclear mature, cytosolic and membrane half-lives with transcripts classified by encoded targeting signals (TS) as in (Zinnall *et al*, 2022). Classification from left to right: cytosol-localized transcripts with no TS (n = 6745), nuclear DNA-encoded mitochondrial proteins (n = 807 with 709 being cytosol-localized), transcripts with tail-anchored transmembrane proteins (n = 86), membrane-localized transcripts with no known TS (n = 560), transcripts encoding signal peptides (n = 303) or transmembrane helices (n = 926) or both (n = 52). Center lines of violin plots depict the median values. Yellow and red colors indicate cytosol and membrane localization, respectively.

Next, we investigated the relationship between subcellular mRNA localization and dynamics. Membrane half-lives are high for membrane-localized transcripts, while cytosolic half-lives are high for cytosol-localized transcripts, being quantitatively similar in both compartments with half-lives around 30 min (see Fig. 4B). This seems to indicate that there is no significant difference of transcript stability between mRNAs that are translated by ribosomes in the cytosol and mRNAs being targeted to and translated at the ER. Once a membrane-associated transcript is in the cytoplasm, it reaches the membrane-bound compartment rather fast, with timescales ranging from less than a minute to around ten minutes. The degree of membrane enrichment influences dynamics, with higher membrane enrichment leading to shorter cytosolic and longer membrane stability. On the other hand, the degree of cytosolic enrichment seems to have little influence on cytosolic stability.

We next examined if transcripts encoding different targeting sequences show different behavior. We observed that the different targeting signals have a similar median cytosolic half-life of around 2 min (see Fig. 4C). Membrane-localized mRNAs encoding no known targeting signal surprisingly show the shortest median cytosolic half-life, while transcripts encoding only a signal peptide tend to have a slightly longer median cytosolic half-life. Distributions of membrane half-life vary slightly for different targeting signals, with transcripts encoding transmembrane helices having the lowest median (21 min) and transcripts encoding proteins with signal peptides having the highest (36 min). mRNA encoding tail-anchored proteins, with the first transmembrane helix close to the C-terminus, have a much longer median cytosolic half-life than other co-translationally targeted transcripts. This suggests that co-translational targeting to the ER for transmembrane helix-containing genes happens after the transmembrane helix is translated. Cytosolic or membrane half-lives did not correlate with the number of encoded transmembrane helices, further suggesting that the distance of the first transmembrane helix to the N-terminus influences cytoplasmic kinetics. Nuclear-encoded mitochondrial transcripts are mostly cytosol-localized (709 out of 807) and tend to have similar kinetics to other cytosol-localized transcripts except for nuclear half-lives, which are significantly longer (median of 150 min). Only 98 mitochondrial transcripts are membrane-localized and have similar cytoplasmic kinetics as ER-localized mRNAs, suggesting that most mitochondrial proteins are post-translationally targeted to mitochondria and only a small fraction are co-translationally targeted to the outer mitochondrial membrane.

### Validation with external datasets and orthogonal approaches

mRNA half-lives have mostly been estimated on whole-cell level. To be able to compare our subcellular half-lives for mESC to those whole-cell half-lives from previous studies, we used our model to predict model-derived whole-cell half-lives from the kinetic parameters and the estimated relative abundance of the three fractions (see Fig. 1D and Methods for details). Herzog et al. estimated whole-cell RNA half-lives in mESCs as a proof-of-concept when establishing SLAM seq (Herzog *et al*, 2017). Our model-derived and Herzog et al. whole-cell half-lives show high correlation (r = 0.80, see Fig. 5A). For transcripts with half-lives longer than 4 h, values agree both by ranking and quantitatively (90th percentile 6.3 h and 6.9 h for model-derived and Herzog et al., resp.). Interestingly, we find a diverging trend for half-lives that are shorter than 2 h: the half-lives derived by Herzog et al. show a high density around 1.5 h, while our model-derived half-lives have a longer tail with values ranging down to 0.5 h (10th percentile 1.1 h and 1.9 h for model-derived and Herzog et al., resp.). This may indicate a lower sensitivity of detecting shorter half-lives for the whole-cell pulse-chase design.

**Figure 5:**
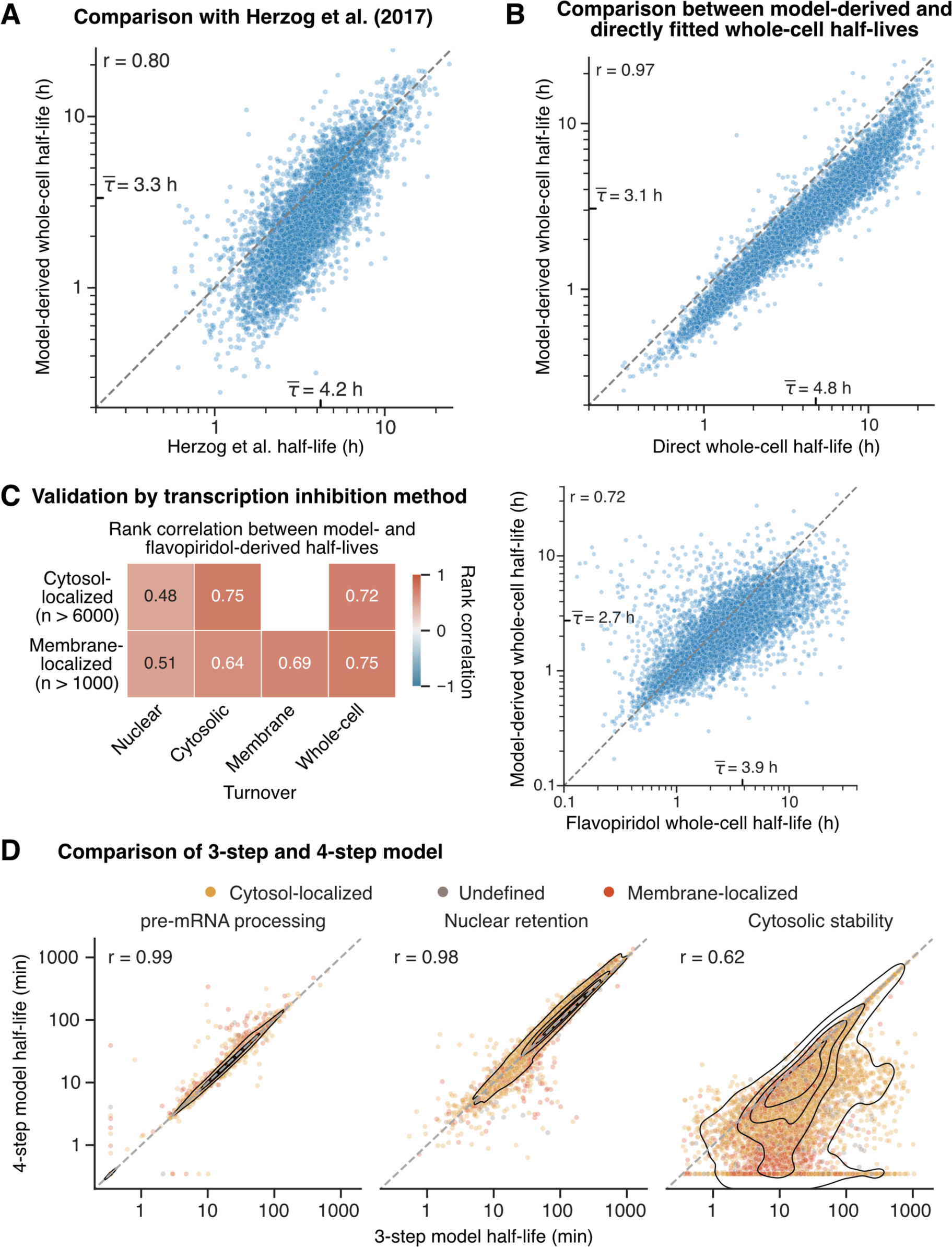
Validation of subcellular parameter estimates with external datasets and orthogonal approaches. **(A)** Comparison with results from first SLAM-seq experiment (Herzog *et al*, 2017). Scatterplot of model-derived (y-axis) and Herzog et al. (x-axis) whole-cell half-lives (n = 5633). Model-derived whole-cell half-lives are calculated from subcellular rates, see Supp. Fig. 4A and Methods. Mean half-life is annotated on the corresponding axis. Spearman rank correlation is shown on top left. Dashed, gray line is the identity line. **(B)** Scatterplot of model-derived and directly-fitted whole-cell half-lives (n = 9860). Model-derived whole-cell half-lives as in (A). Direct whole-cell half-lives obtained by fitting a simple “one minus exponential decay” model to the share of new to total mRNA from whole-cell extract data. Mean half-life is annotated on the corresponding axis. Spearman rank correlation is shown on top left. Dashed, gray line is the identity line. **(C)** Comparison of model-to flavopiridol-derived half-lives. Left: Correlation heatmap of model-with flavopiridol-derived subcellular half-lives split into cytosol-(upper row) and membrane-localized genes (lower row). Values in heatmap are spearman rank correlations, also indicated by color. For details on how half-lives using flavopiridol were derived, see Methods and Supp. Fig. 5. Right: Scatterplot of model-derived and flavopiridol whole-cell half-lives (n = 8385). Model-derived whole-cell half-lives as in (A). Mean half-life is annotated on the corresponding axis. Spearman rank correlation is shown on top left. Dashed, gray line is the identity line. **(D)** Comparison of 3-step and 4-step model, which model mRNA flow until cytosolic and membrane-bound compartments, respectively. Scatterplot of nuclear pre, nuclear mature and cytosolic half-lives with values from 3-step and 4-step model on x- and y-axis, respectively (n = 8165, all genes were fit with both models). Black lines are 2-dimensional KDE to indicate density of points. For details on the models, see Model sections in Results and Methods. Spearman rank correlation is shown on top left. Dashed, gray line is the identity line.

In order to augment the robustness of our investigation into the estimated half-lives, we further compared our model-derived half-lives with a simple “one minus exponential decay” model that we fit on SLAM-seq data of our whole-cell samples, referred to as “direct” whole-cell half-lives in the following. Comparing the direct to the model-derived whole-cell half-lives, we find a near-perfect rank correlation (r = 0.97), but the values were systematically lower in our compartment model (see Fig. 5B; mean half-life 3.1 h and 4.8 h for model-derived and direct, resp.). This bias was most likely a result of the length-dependent labeling bias as well as the non-exponential nature of the process. Interestingly, the 90th percentile value of the direct half-lives is higher compared to both our model-derived half-lives and those estimated by Herzog et al. (10.4 h, 6.3 h and 6.9 h, resp.).

To experimentally validate our metabolic labeling-derived subcellular flow rates with an orthogonal (but more interfering) method, we inhibited transcription using flavopiridol (Chao & Price, 2001) followed by subcellular fractionation and RNA sequencing, resulting in a second set of subcellular rates (see Methods for details). Reassuringly, we find high correlation between the two sets of cytosolic, membrane and whole-cell half-lives for both cytosol- and membrane-localized mRNAs. Interestingly, the correlation was only moderate for nuclear mature half-lives (see Fig. 5C left) which may be due to stress-induced changes in nuclear RNA turnover. When comparing the flavopiridol- and metabolic labeling-derived whole-cell half-lives (Fig. 5C, right), we find that for half-lives longer than 4 h, flavopiridol-derived half-lives were generally longer than metabolic labeling-derived ones, suggesting reduction in mRNA decay after transcription blockage.

As we used different models for cytosol and membrane-localized mRNAs (3 step and 4 step model, respectively), we asked how model choice influences the kinetic parameters. We therefore fitted both mathematical models to our subcellular SLAM-seq data, and compared the half-lives between the two models. We found a near-perfect correlation for pre-mRNA processing (r = 0.99) and nuclear retention (r = 0.98), which are modeled the same way in both models (see Fig. 5D). However, for the cytosolic stability we find lower agreement (r = 0.62), as here the 4-step model used additional information to constrain the ratio between cytosolic and membrane half-lives. This results in all membrane-localized, but also many cytosol-localized transcripts exhibiting lower cytosolic half-lives in the 4-step than in the 3-step model. For the former, this was desired, as T2C mutations are distinct between cytosol and membrane compartments, whereas for the latter, this is undesirable, as the T2C mutations are similarly frequent in both membrane and cytosol compartments and the fit then systematically underestimates the cytosolic half-life to accurately fit the membrane half-life.

Taken together, our validation efforts using our and published data sets suggest that the compartment-derived rates provide a high-quality quantitative assessment of mRNA turnover.

## Discussion

Our study provides a comprehensive analysis of and resource for nucleocytoplasmic mRNA kinetics in mESCs. Applying mathematical modeling to time-resolved subcellular SLAM-seq data we were able to quantify intracellular mRNA flow and offer valuable insight into the dynamics of the mRNA metabolism. Previously, mRNA kinetics were studied mostly on whole-cell level, while our approach yields information on subcellular kinetics, even dissecting cytoplasmic into cytosolic and membrane dynamics, distinguishing it from recent studies on subcellular mRNA kinetics (preprint: Smalec *et al*, 2022; preprint: Müller *et al*, 2023).

Here, we modeled subcellular mRNA kinetics using a linear, inhomogeneous system of ordinary differential equations, accounting for different steps in the life cycle of mRNA molecules. Incorporating steady-state ratios in the fitting procedure allowed us to distinguish between export and degradation in the nuclear compartment and ensured that the fitted parameters align with the measured subcellular mRNA expression levels. The quantitative accuracy of the steady-state ratios was ensured by accounting for the varying amounts of mRNA present in the different cellular compartments in a manner similar to a recently published method (Dai *et al*, 2022).

To account for an observed labeling bias towards the 3’ end of transcripts when using poly(A)-selection and full-length transcript sequencing, we included a transcript elongation rate in our model. This modeling approach allowed for the estimation of kinetic parameters of pre-mRNA processing, nuclear retention, cytosolic and membrane stability for mRNAs on a global level, including an approximation of transcriptional elongation rates. Our estimated mean elongation rates of 1.2 to 2.6 kb/min for shorter and longer genes, respectively, are highly similar to median elongation rate estimations of 1.25 to 1.75 kb/min measured before in five different human cell lines (Veloso *et al*, 2014). Furthermore, our analysis revealed that transcriptional elongation was higher for longer genes, as observed previously (Veloso *et al*, 2014; Jonkers *et al*, 2014). In both of these two studies, differences in gene length-specific elongation rates were correlated with distinct histone modifications, suggesting that gene structure and epigenetic modification influence RNA polymerase II elongation rates.

Across transcripts, the timescales of the subcellular kinetic parameters span multiple orders of magnitude. One of the key findings of our study was the observation that nuclear retention is the rate-limiting step for most transcripts, as evidenced by the significantly longer mean half-lives of 96 min for mature mRNA in the nucleus compared to the cytoplasm half-life of 19 min. This finding is consistent with prior works applying imaging-based approaches, demonstrating that nuclear retention of transcripts serves as an effective mechanism for buffering noise (Battich *et al*, 2015; Bahar Halpern *et al*, 2015). Battich and coworkers derived nuclear retention times between ∼5–90 min for close to 300 newly synthesized transcripts, with a median of ∼20 min (Battich *et al*, 2015). However, the authors argued that these nuclear retention times are likely an underestimation for most other genes, since these genes were fast-responding genes during stress signaling. Moreover, Halpern and colleagues showed that in mouse tissues spliced and polyadenylated mRNAs are retained in the nucleus for many protein-coding genes to reduce cytoplasmic gene expression noise (Bahar Halpern *et al*, 2015). Ren and coworkers found that a substantial fraction of newly synthesized transcripts was retained in the nucleus even after 6 h, corroborating the previous observation of nuclear mRNA retention (Ren *et al*, 2023). Similarly, by applying sequencing of metabolically labeled RNA in the nuclear and cytosolic compartments followed by mathematical modeling, Mueller et al. recently reported that mRNA molecules generally spend most of their life in the nucleus, highlighting nuclear retention as a critical determinant of subcellular mRNA dynamics (preprint: Müller *et al*, 2023). The relative amount of poly(A) mRNA in the nucleus varies across cell types (Bahar Halpern *et al*, 2015; Dai *et al*, 2022; Gondran *et al*, 1999). As the nucleus makes up a large part of the cell volume in mESCs and we observe a high relative nuclear mRNA amount, the nuclear retention half-lives estimated here are likely to be higher than in other, differentiated cell types.

To identify functional categories associated with genes exhibiting extreme subcellular rates, we undertook gene set enrichment analyses. mRNAs transcribed from genes involved in transcription regulation were enriched among those with short nuclear half-lives. Conversely, mRNAs from genes associated with translation and metabolic processes, in particular mitochondrial and ribosomal genes showed enrichment among those with longer nuclear half-lives. Interestingly, mRNAs of nuclear-encoded mitochondrial genes presented mostly similar cytoplasmic kinetics when compared to cytosol-localized transcripts, suggesting mostly post-translational targeting to the mitochondria with only around 10% of transcripts being co-translationally targeted.

Using cytosolic and membrane mRNA expression levels, we were able to identify cytosol– and membrane-localized transcripts with the latter mostly being associated with the ER membrane. These membrane transcripts were localized within two minutes to the ER, mostly independent of the targeting signal, and then were stable for around 30 min at the ER, with transcripts targeted by the signal recognition particle having slightly longer half-lives. Cytosolic half-lives of cytosol-localized transcripts were centered around half an hour, revealing that cytosol- and membrane-localized mRNAs have similar cytoplasmic stability, suggesting that differences in protein expression between cytosol- and membrane-localized transcripts with similar mRNA levels derive from differences in translational efficiency (Voigt *et al*, 2017; Lashkevich & Dmitriev, 2021; Zinnall *et al*, 2022). An intriguing prospect for future inquiry revolves around the impact of translation on the subcellular localization of mRNA. Recent findings have shown that, notably, when translation initiation is inhibited, the ER becomes the predominant site for the localization of newly exported mRNAs, prompting questions about the intricate relationship between translation and subcellular mRNA localization (Child *et al*, 2023).

Since nuclear retention is the rate-limiting factor, it is harder to correctly estimate all processes taking place afterwards, especially if they happen on a comparatively short time scale. Using the well-measured steady-state ratios helps to essentially determine the short and difficult to estimate cytosolic stability from the longer and more easily determined membrane stability in the case of membrane-localized transcripts, where slight contaminations between nuclear and membrane compartment would further hinder correct estimation.

Based on subcellular parameters we calculated model-derived whole-cell half-lives that correlate highly with previous experimentally determined whole-cell half-life estimates. This analysis provided confidence in the accuracy of the subcellular rates and their ability to represent whole-cell mRNA kinetics. Both model-derived and direct whole-cell half-lives were correlated highly with nuclear retention half-lives. To validate the subcellular flow rates further, we inhibited transcription and performed subcellular fractionation, generating a second set of subcellular rates. These rates showed good agreement with the original rates, further supporting the reliability of the findings.

In conclusion, our findings offer deeper insights into the dynamics of mRNA metabolism and uncover compartment-specific features of post-transcriptional regulation, showing that mature mRNAs spend most of their lifetime in the nucleus in mESCs. The relationship between mRNA kinetics and gene functions give directions for further investigation to analyze the RNA-bindng proteins and molecular processes, like translation, influencing subcellular mRNA dynamics. These findings provide a foundation for future research into the mechanisms of mRNA processing and localization within mammalian cells, ultimately contributing to our broader understanding of gene expression regulation in a subcellular context.

## Materials and Methods

### Mouse embryonic stem cell (mESC) culture

E14TG2a mESCs (Iacovino *et al*, 2014) were cultured in 0.1% gelatin [w/v] coated plates in “2i + LIF” ES medium [Advanced DMEM/F12 (12634028, Thermo Fisher Scientific) – Neurobasal (21103049, Thermo Fisher Scientific) – Knockout™ DMEM (10829018, Thermo Fisher Scientific) (1 : 1 : 0.5), 14% Fetal Bovine Serum qualified for ES cells (16141079, Life Technologies), 1X N2 (17502048, Thermo Fisher Scientific), 1X B27 (17504001, Thermo Fisher Scientific), 1X GlutaMax (35050061, ThermoFisher Scientific), 1X MEM Non-Essential Amino Acid (11140050, Thermo Fisher Scientific), 1X Nucleosides (ES-008-D, Merck Millipore), 100 µM β-mercaptoethanol (21985023, Thermo Fisher Scientific), 3 μM CHIR99021 (SML1046-5MG, Invitrogen) and 1 μM PD0325901 (PZ0162-5MG, Invitrogen), 1000 U/ml Leukemia inhibitory factor (ESG1107, Merck Millipore)], under a controlled atmosphere at 5% CO2 and 37°C. mESCs were seeded the day before the experiments at a density of 3× 105 cells/ml.

### Metabolic labeling and cell fractionation

Two independently passaged biological replicates of mESCs (∼3.5 × 107 cells per replicate) were separately labeled in “2i + LIF” ES medium supplemented with 500 µM (for 0, 15, 20, 30, 40 min) or 100 µM (for 60, 120 and 180 min) 4-thiouridine (4sU, RP-2304, ChemGenes) and fractionated by sequential detergent extraction, as described previously (Jagannathan *et al*, 2011) with minor modifications. Briefly, medium was aspirated, 5 ml of ice-cold PBS supplemented with 100 µM cycloheximide (A0879,0001, Biochemika) was added to the plates, cells were scraped from the plates, transferred to 15ml falcon and spun down. Pellet was resuspended in 500 µl of ice-cold permeabilization buffer (110 mM KOAc, 25 mM K-HEPES pH 7.2, 2.5 mM Mg(OAc)2, 1 mM EGTA with freshly added 0.015% digitonin, 1 mM DTT, 100 μg/ml cycloheximide, 1X Complete Protease Inhibitor Cocktail and 40 U/mL RNaseOUT™). 100 µl of the sample was taken aside as Total extract and rest was incubated for 10 min at 4°C with rotation, followed by centrifugation at 3000 g 5 min at 4°C. Supernatant (corresponding to the Cytosolic fraction) was transferred to new tube while the pellet was resuspended in 5ml of wash buffer (110 mM KOAc, 25 mM K-HEPES pH 7.2, 2.5 mM Mg(OAc)2, 1 mM EGTA with freshly added 0.004% digitonin, 1mM DTT, 100μg/ml cycloheximide) and spun down again at 3000 g 5 min at 4°C. After centrifugation washed pellet was mixed with 500 µl of ice-cold lysis buffer (400 mM KOAc, 25 mM K-HEPES pH 7.2, 15 mM Mg(OAc)2, 0.5% (v/v) NP-40 and freshly added 1 mM DTT, 100 μg/ml cycloheximide, 1X Complete Protease Inhibitor Cocktail, 40 U/mL RNase Out) and incubated for 5 min on ice followed by centrifugation at 3000 g 5 min at 4°C to collect the supernatant (corresponding to the Membrane fraction) and the pellet (insoluble and the nuclear fraction). For additional purity nuclei were loaded on 10% sucrose cushion in lysis buffer and centrifuged at 200 g 5 min at 4°C. The cytosolic and membrane fractions were clarified at 7500 g 10 min at 4°C to remove cell debris. 20 µl of all fractions were taken for Western analysis while the rest were mixed with Trizol LS (10296028, Thermo Fisher Scientific) for subsequent RNA isolation.

### Western blotting

Protein lysates from cell fractionation experiments were separated on 10% SDS PAGE, transferred to nitrocellulose blotting membrane (10600004, GE Healthcare), blocked in 5% dry milk and probed for cytosolic (beta-tubulin, GAPDH), membrane (BCAP31), nuclear (Lamin A/C, TBP, H3) markers and S6 ribosomal protein. Antibodies were used at a dilution of 1:5000 for anti-beta-tubulin (T8328-200UL, Sigma-Aldrich, mouse), 1:25000 for anti-GAPDH (G8795, Sigma-Aldrich, mouse), 1:2000 for anti-TBP (ab818, Abcam, mouse), 1:500 for anti-Lamin A/C (14-9688-80, Invitrogen, mouse), 1:1000 for anti-H3 (ab1791, Abcam, rabbit), anti-BCAP31 (11200-1-AP, Proteintech, rabbit) and anti-S6 (2217, Cell Signaling Technology, rabbit) and detected by 1:2000 dilution of respective secondary HRP-antibody-conjugates (anti-rabbit, P044801-2, Agilent; anti-mouse, P044701-2, Dako).

Primary antibodies were incubated at room temperature for one hour followed by 3 washing steps for 5 min and incubation with secondary antibodies for one hour. Images were acquired using Amersham ECL Western Blotting Detection Reagent (RPN2209, GE Healthcare) on an Amersham Imager 680 (GE Healthcare).

Transcriptional inhibition by flavopiridol

Three independently passaged biological replicates of mESCs (∼3.5 × 107 cells per replicate) were cultured in “2i + LIF” ES medium supplemented with 1µM flavopiridol (HY-10005-10mg, Biotrend) to block the transcription. Cells were harvested at 0, 30, 60, 120 and 180 min after addition of flavopiridol followed by cell fractionation as described above. Obtained fractions were mixed with Trizol LS and RNA was isolated following the manufacturer instructions. One microgram of total RNA was used as an input for TruSeq Stranded mRNA Library Prep Kit according to the instructions of the manufacturer. The multiplexed libraries were sequenced using HiSeq 4000 for pair-end 75 cycles by the BIH Genomics platform at the Max Delbrück Center for Molecular Medicine.

### RT-qPCR

For assessment of stress response caused by 4sU toxicity mESCs were incubated with 100, 250, 500 µM 4sU for 0, 60, 120 and 240 min with subsequent RNA isolation. Reverse transcription was performed by SuperScript III Reverse Transcriptase (18080085, Thermo Fisher Scientific) and random hexamers (48-190-011, Thermo Fisher Scientific) following the manufacturer’s instruction. Around 100 ng of the synthesized cDNA was used as an input for 20 μl qPCR reaction using SYBR Green PCR Master Mix (Applied Biosystems) with the gene specific primer pairs targeting stress-responsive gene p21 (forward: 5’-TCGCTGTCTTGCACTCTGGTGT-3’, reverse: 5’-CCAATCTGCGCTTGGAGTGATAG-3’). The mean CT value was calculated for three biological replicates.

### Bioinformatics and data analysis

Alignment of raw reads was done with STAR (v2.7.6). Reads in exons were counted with HTSeq-count (v0.11.1) and TPM values were obtained by RSEM (v.1.3.3) using default parameters. Genes were defined as expressed if they had an average TPM value ≥1.5 in at least one subcellular fraction. Membrane-to-cytosol enrichment was equal to the fold change between the average TPM values of membrane and cytosolic samples. To define membrane-bound and cytosolic mRNAs we used membrane enrichment cutoffs ≥3.0 and ≤1.5, respectively, for all expressed mRNAs. T counts and T2C mismatches in raw reads were counted and mapped to exons/introns using a custom script. Based on a binomial mixture model by Juerges et al. (2018) we determined the conversion rate per sample (Supp. Fig. 1C). With the conversion rates, T and T2C conversion counts the share of new to total RNA was calculated on intron/exon level, which was used later on as input for the kinetic model (see Figs. 1A bottom and 1F). Fitting was performed throughout with the python package lmfit (v1.0.3) using the Levenberg-Marquardt “least-squares” algorithm to minimize the chi-squared unless otherwise specified. Gene set enrichment analyses were performed with fgsea (v1.28.0) with gene set sizes limited from 100 to 600. For general data analysis purposes, python 3.9 and R v4.3.2 were used. Read mapping, counting, T2C data processing and model fitting was implemented in Snakemake (Mölder *et al*, 2021).

### Minimal effect of metabolic labeling on the transcriptome

To assess the effect of the metabolic labeling on transcriptional output, we performed a differential gene expression analysis on RNA count data from whole-cell extracts using DESeq2 (v1.40.2). Comparing all subsequent time points to the t=0 time point, we found only three differentially expressed genes at 180 min and otherwise none (see Supp. Fig. 1A, log2FC > 1 and adjusted p-value < 0.1). This shows a negligible effect of 4sU labeling on the transcriptome.

### Quantifying relative mRNA abundance of subcellular compartments

To get a biologically accurate quantification of the RNA expression ratio between different subcellular compartments, we need to quantify how much RNA is present in those compartments. This allows us to constrain the parameter space by steady-state expression ratios and to reconstruct whole-cell dynamics from subcellular kinetic parameters. Under the assumption that, in the gene space containing all expressed genes, the RNA expression vector of the whole-cell extract can be approximated by summing the RNA expression vectors of the three subcellular fractions with a corresponding factor for each, we fitted the model *a* · TPM_nuc_ + *b* · TPM_cyto_ + *c* · TPM_mem_ = 1 · TPM_whole-cell_ to the whole-cell RNA expression with the constraint *a + b + c* = 1. RNA expression vectors were the mean over all replicates and time points of the corresponding fraction (subcellular or whole-cell) in gene space. The inverses of the standard errors were used as weights in the fitting procedure. This resulted in the following values for the relative abundances: nuclear factor = 0.33, cytosolic factor = 0.20 and membrane factor = 0.47 (see Fig. 1D). It is of note that the cytosolic abundance is the lowest. The subcellular TPM counts are multiplied by their corresponding factor before calculating the steady-state ratios that are used to constrain the parameter space in the fits (Box 1).

### Model fitting of subcellular mRNA dynamics

Summarizing different models and genes, the following kinetic rates were estimated: transcript elongation rate, pre-mRNA processing rate, nuclear export rate, nuclear decay rate, cytosolic decay rate, cytosolic transport rate and membrane decay rate (see Box 1). The half-lives, calculated from those rates, were primarily shown in the results section. Per gene, the analytical solutions of the rescaled system, shown in Box 2, were used to fit all corresponding kinetic rates simultaneously to the subcellular, time-resolved labeling data. More specifically, the mean share of new to total RNA for exons and introns across replicates was used as input data for the fit (see example gene with data from cytosolic compartment in Fig. 2B). Each data point was weighted by the inverse of a mixed standard error across replicates, with the mixed standard error consisting half of the standard error per time and compartment and half of the standard error averaged over time per compartment. Acting as a variance-stabilization mechanism, the minimum of the standard error for each exon was set to 0.06 (the mean standard error across all exons at t=15 min). As parameter boundaries a minimum of 0.001 min^−1^ and a maximum of 2 min^−1^ was set for all rates (for decay rates the minimum was set to 1e-06 min^−1^). To further constrain the parameter space we used the steady-state expression ratios via the equations shown in Box 1 and the “expr” keyword from lmfit. The upper and lower limits between parameter ratios were given by the steady-state ratios plus and minus five times its standard error. Per gene, the kinetic parameters were fit in logarithmic space 200 times with random initial conditions using a computationally inexpensive local fitting method. Out of the ten lowest chi-squared fits, the result with parameter values farthest from the parameter boundaries was chosen as best and final fit. The quality control involved three steps: genes were excluded if the best fit showed a) reduced chi-squared > 4, b) nuclear or cytosolic (cytosol-localized) or nuclear and membrane (membrane-localized) parameter values at the maximum allowed boundary, c) relative standard deviation (of the ten least chi-squared fits) of any parameter > 0.00001 (see Supp. Figs. 2A, 2B and 2C).

#### Box 2.

##### Analytical solutions of the rescaled system (see Box 1) and their usage in the fitting procedure

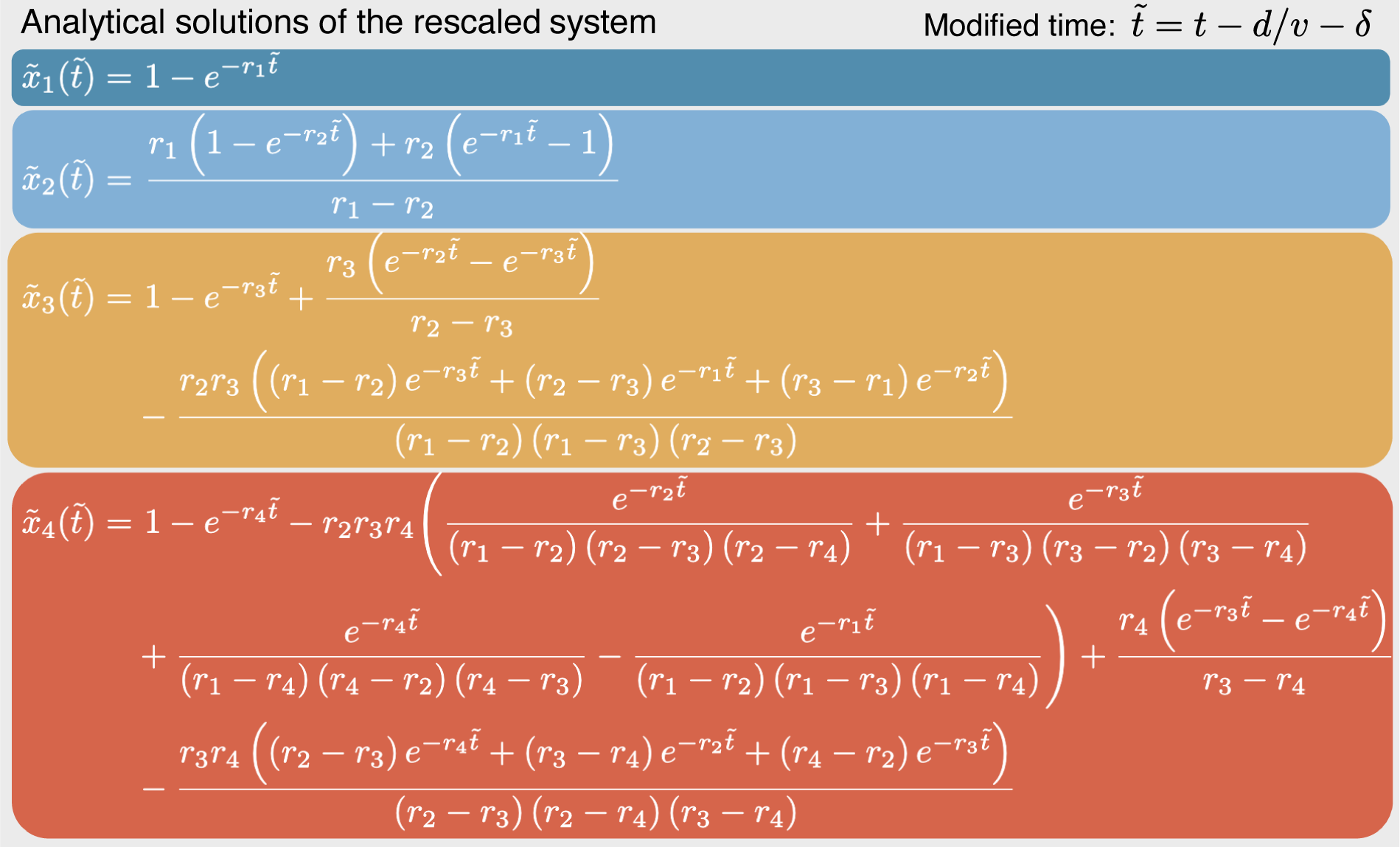

For intron-containing genes (n = 8548), solutions of the first, second, third and fourth step are used to fit nuclear pre-, nuclear mature, cytosolic and membrane-bound mRNA, respectively, with the fourth step being used only for membrane-localized and undefined genes (n = 1726). As indicated on the top right, we incorporate the transcript elongation rate *v* to time-delay the solutions: for mature mRNA we fit on exon-level and *d* is the distance of the exon to the 3’ end, and for pre-mRNA we fit all intronic regions together and *d* is the gene length times 0.1, with this heuristic factor being the mean relative exon distance to 3’ end across genes weighted by expression. *δ* is the overall delay of 5 minuntil any labeling is experimentally observed. The modified time is set to 0 for all negative values. For intron-less genes or genes with reliable T2C data present only for one exon (n = 1314), the first, second and third solutions are used to fit nuclear mature, cytosolic and membrane-bound mRNA, respectively, with the third step being used only for membrane-localized or undefined genes (n = 318). In this case transcript elongation and pre-mRNA processing rates are not fit and *t̃* = *t* – *δ*.

### Nuclear mRNA degradation contributes to mRNA dynamics for 40% of fitted genes

The role and contribution of nuclear decay of poly(A) mRNA has been under discussion (Schmid & Jensen, 2018). To test the contribution of nuclear decay in subcellular mRNA dynamics, we included nuclear degradation by default in our model. While more than half of the fitted transcripts show no nuclear degradation, for roughly 4000 transcripts nuclear degradation contributes to the overall nuclear dynamics of mature mRNA (see Supp. Fig. 4A). Ranking genes by the share of nuclear degradation at overall nuclear kinetics and performing a GSEA showed that transcripts encoding membrane, ribosomal and mitochondrial proteins were upregulated, so tend to have high nuclear degradation, while transcription factors were strongly down-regulated (see Supp. Fig. 4B). We then investigated the dynamics of all ribosomal and nuclear-encoded mitochondrial proteins and found that in both groups only a subset of transcripts showed substantial nuclear decay with their distributions even being similar to the rest of transcripts (see Supp. Fig. 4C). Surprisingly, also a small share of transcription factors showed nuclear degradation. This suggests that nuclear degradation is more closely linked to sequence-specific molecular properties than to biological function. However, as we modeled nuclear degradation by default for all transcripts, further improvements on the modeling procedure would help to more reliably determine the nuclear decay parameter, such as modeling with and without the parameter and using a Bayesian information criterion to decide per transcript if the parameter should be included.

### Transcription inhibition using flavopiridol

To validate our metabolic labeling-derived subcellular flow rates, we inhibited transcription in mESCs using flavopiridol for 0, 30, 60, 120 and 240 minutes followed by subcellular fractionation and RNA sequencing (as specified above). Reads were aligned with STAR (v2.7.6) and counted with RSEM (v1.3.3). Library sizes were normalized using genes with known half-lives greater than 14 hours (Herzog *et al*, 2017). Samples of each time series were further normalized to the corresponding t=0 time point (Supp. Fig. 5C). We fitted a simple exponential decay model per compartment for roughly 10,000 genes, giving the aggregated subcellular turnover rates (see example in Supp. Fig. 5D), in contrast to the rates of transition from one compartment to the next. Transcription inhibition-derived half-lives are shown in Fig. 5C and Supp. Fig. 5E.

### Model-derived (whole-cell) half-lives

To compare the transitional, metabolic labeling-derived half-lives to the aggregated, flavopiridol-derived half-lives, we generated model-derived pulse-chase data from the transitional rates using the equations in Box 2 and the fact that the share of old mRNA is given by one minus the share of new mRNA. To the model-derived pulse-chase data (n = 9862) we fitted a simple exponential decay model (see example fit for a single gene in Supp. Fig. 5A). For the subcellular compartments the equations in Box 2 can be used as is, while for the whole-cell compartment the subcellular solutions are summed with their corresponding relative RNA abundance factors. Model-derived aggregated half-lives (subcellular and whole-cell) are used in Supp. Fig. 5B and Fig. 5E. Model-derived whole-cell half-lives are also used in Fig. 5A.

### Targeting signal and gene group annotations

Signal peptide and transmembrane helix annotations were downloaded from Ensembl Biomart. We defined tail-anchored proteins as those transmembrane domain-containing proteins lacking signal peptide, for which the first transmembrane helix started 50 or less amino acids from the C-terminus. A list of mitochondrial proteins encoded in the nuclear DNA was obtained from Mitocarta 2. Ribosomal proteins were defined as those that contained an “Rps” or “Rpl” prefix in their gene symbol. Transcription factors were defined by a list of mESC transcription factors from the <u>Gifford lab</u>.

## Data and code availability

Metabolic labeling RNA-seq data are accessible from the NCBI Gene Expression Omnibus (GEO) under the accession number GSE252199. Flavopiridol transcription inhibition RNA-seq data are accessible from the NCBI Gene Expression Omnibus (GEO) under the accession number GSE256335. Data processing and model fitting scripts are publicly available on GitHub at https://github.com/steinbrecht/subcellular-SLAM.

## Acknowledgements

We thank Stefan L Ameres for help with the experimental design. We acknowledge the Genomics technology platform of the Max Delbrück Center for Molecular Medicine for high-throughput sequencing. Computation has been performed on the HPC for Research cluster of the Berlin Institute of Health. We thank members of the Blüthgen and Landthaler labs for fruitful discussions. Funding was provided by Deutsche Forschungsgemeinschaft (grants RTG2424 CompCancer and SFB/TRR 186 to D.S.).

## Author contributions

M.L. and N.B. conceived the study. M.L. and I.M. designed the experiments. I.M. carried out all of the experimental work. M.M. and J.M. performed bioinformatic data analysis. D.S. and J.M. performed modeling. D.S. prepared the manuscript and the majority of figures. I.M., N.B., M.L. and M.M. critically revised and contributed to the manuscript.

## Supplementary Figures

**Supp. Fig. 1:**
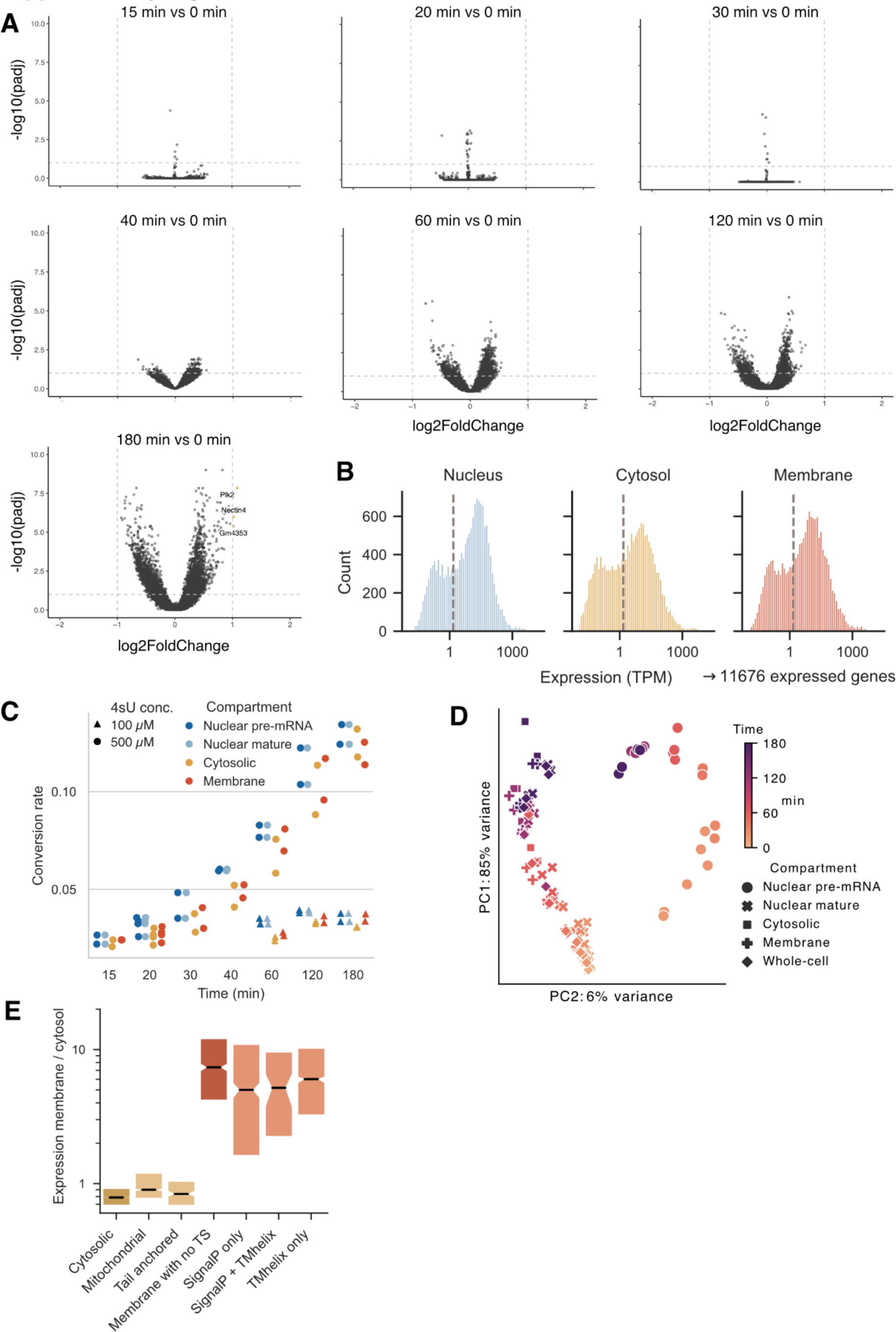
Overview of mRNA expression and T2C labeling data. (A) Differential expression analysis of whole-cell extract RNA sequencing data. Volcano plot showing Benjamini-Hochberg-adjusted p-values against fold changes. Facets show subsequent 4sU labeling times against t=0 point without labeling. Vertical dashed lines indicate a two–fold change in expression. Horizontal dashed lines indicate adjusted p-value of 0.1. Only three genes (Plk2, Nectin4 and Gm4353) are differentially expressed for the 180 min time point. (B) Gene expression in subcellular fractions. Histograms show TPM values averaged over all time points and replicates. Vertical dashed line indicates average TPM of 1.5. If a gene has an average TPM > 1.5 in at least one subcellular fraction, it is considered expressed. (C) Estimates of T2C conversion rate. Conversion rates of individual samples were estimated fitting a Binomial mixture model to T and T2C count data. Color indicates compartment. Shape indicates 4su concentration. (D) Principal component analysis of the T2C labeling data using 2,000 most variable genes. Samples cluster by exposure time to 4sU rather than replicate, with nuclear pre-mRNA being separated from other compartments. (E) Box plot of ratio between membrane and cytosolic expression (membrane enrichment) with transcripts classified by encoded TS as in Fig. 4C. Center lines of box plots depict the median values. Notches show 95% confidence interval of median values. Lower and upper hinges of box plots correspond to the 25th and 75th percentiles, respectively. Yellow and red colors indicate cytosol and membrane localization, respectively.

**Supp. Fig. 2:**
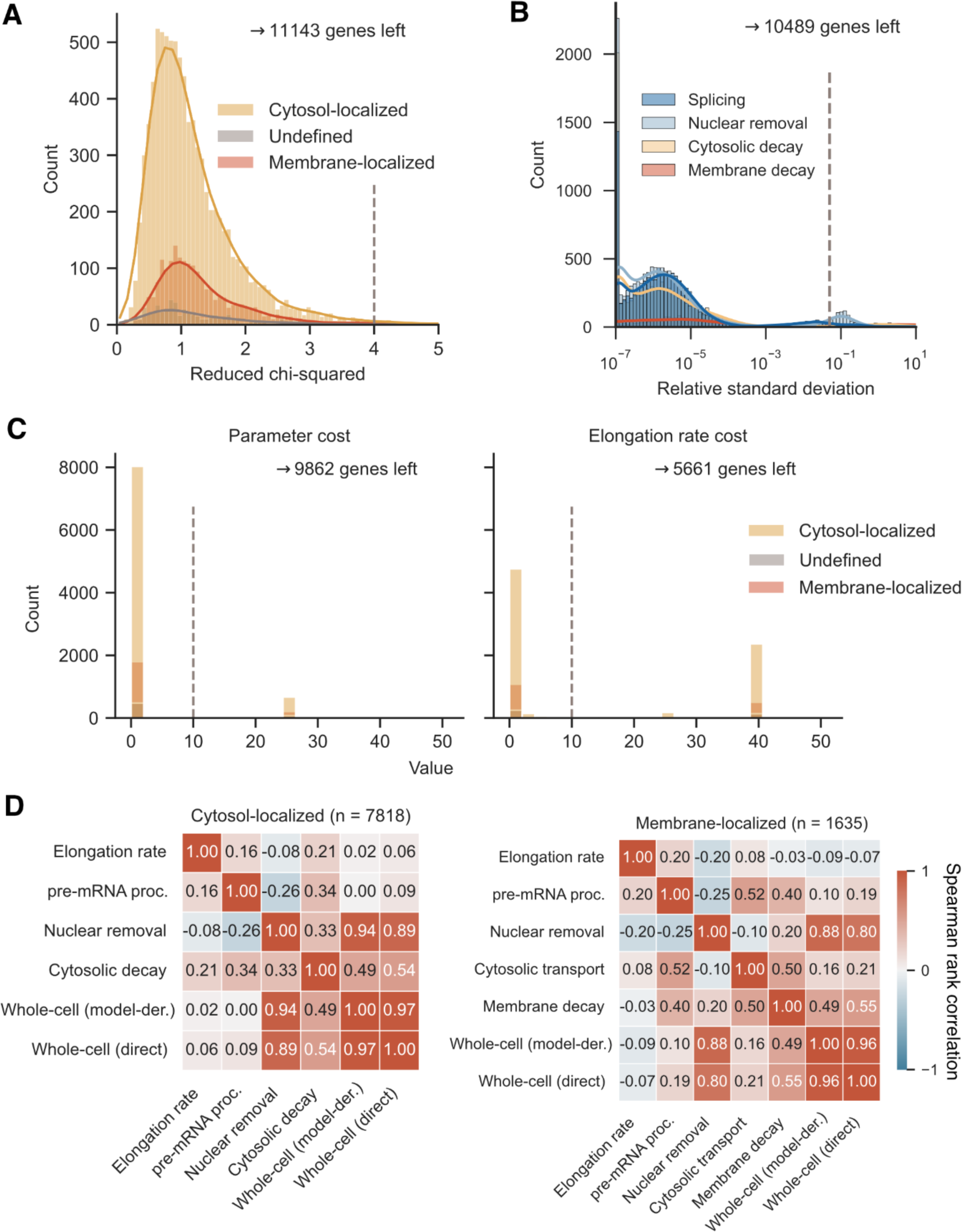
Quality control of fit results. (A) First quality control step. Histograms of reduced chi-squared, which is minimized during fitting (optimal value is 1). Vertical dashed line shows cutoff of 4. If the best fit for a gene has a value higher than the cutoff, the gene is excluded. Colors indicate mRNA localization. (B) Second quality control step. Histograms of relative standard deviation, calculated by dividing the standard deviation of the ten best fit results by the value of the best fit. Vertical dashed line shows cutoff of 0.05. If the relative standard deviation for a gene has a value higher than the cutoff, the gene is excluded. Colors indicate parameter. (C) Third quality control step. Histograms of the boundary cost for all kinetic parameters (left) and elongation rate (right). Boundary cost is near 0 if fit value is far away from upper and lower limits and increases drastically as the value approaches the allowed limits. A high boundary cost therefore indicates that the fit was stuck at maximum or minimum allowed value. Vertical dashed line shows cutoff of 10. If the best fit for a gene has a parameter cost higher than the cutoff, the gene is excluded. Genes with a elongation rate cost higher than the cutoff are only excluded for analyses specifically of the elongation rate. Colors indicate mRNA localization. (D) Heatmaps showing the Spearman rank correlation between kinetic parameters for cytosol-(left) and membrane-localized transcripts.

**Supp. Fig. 3:**
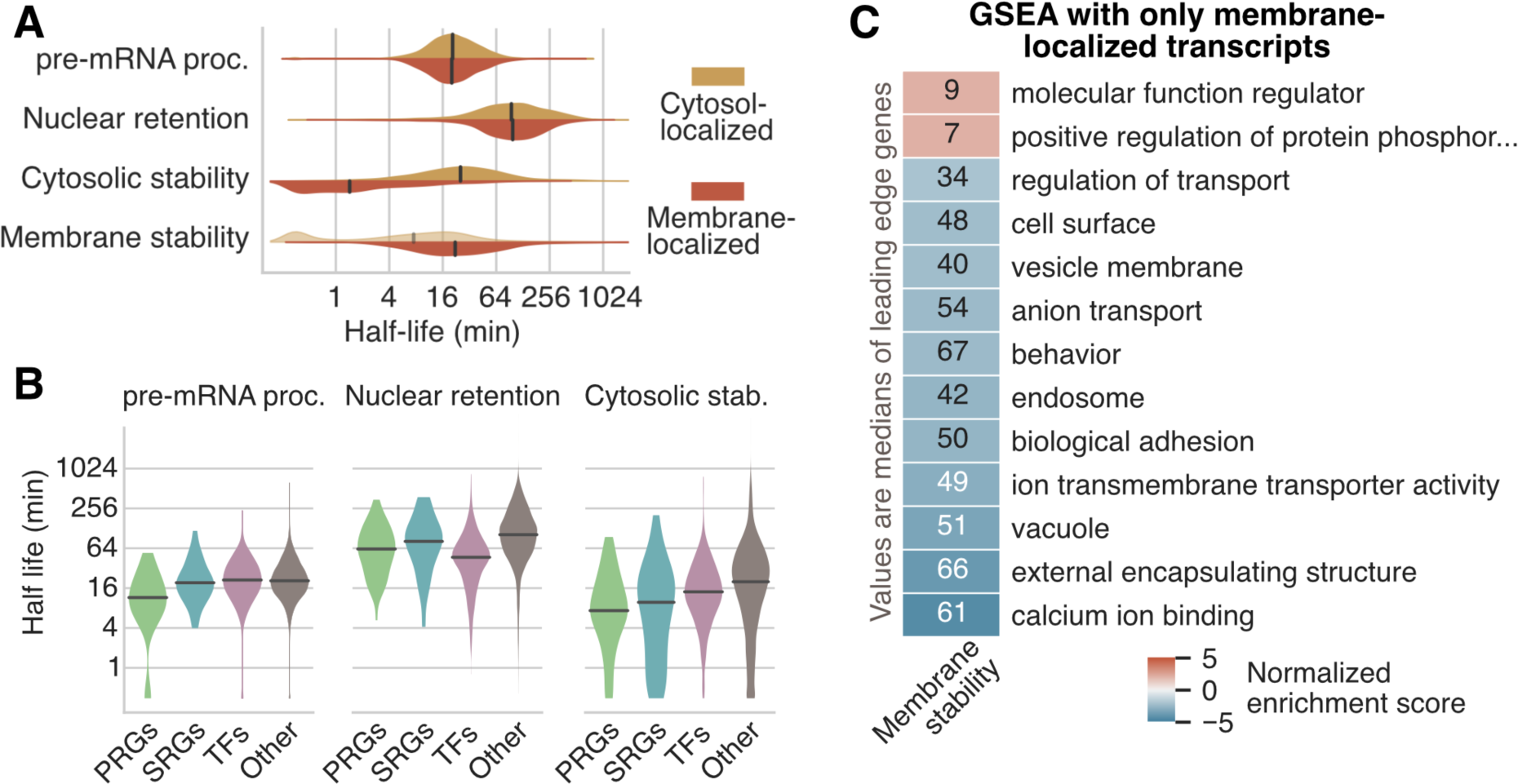
Kinetic differences between different gene groups. (A) Violin plots of pre mRNA processing, nuclear retention, cytosolic, membrane half-lives for cytosol-(n=7818) and membrane-localized (n=1635) transcripts. (B) Violin plots of pre mRNA processing, nuclear retention and cytosolic half-lives for primary response genes (PRGs, n=), secondary response genes (SRGs, n=), transcription factors (TFs, n=) and all other (n=) transcripts. Definitions of PRGs and SRGs are taken from Uhlitz et al. (2017). For the definition of TFs, see Methods. (C) GSEA based on the GO using only membrane-localized transcripts and ranking by membrane decay. All significant terms (FDR < 0.05) are shown. Color indicates normalized enrichment score. Annotated values are the median half-lives of leading edge genes of each term.

**Supp. Fig. 4:**
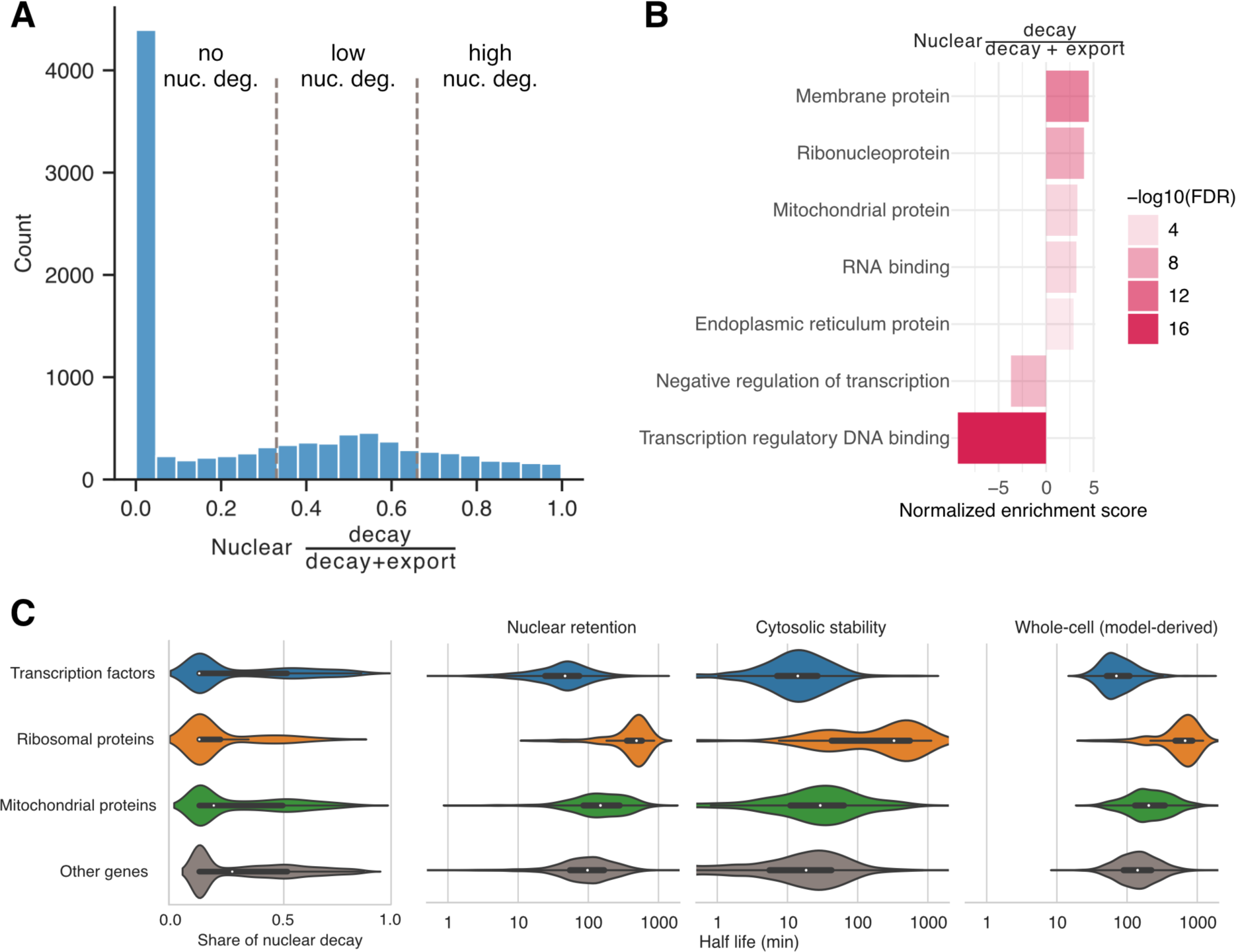
Contribution of nuclear decay to nuclear dynamics. (A) Classification of share of degradation at overall nuclear kinetics. Histogram including kernel-density estimation of ratio between nuclear degradation rate and sum of nuclear degradation and export rates. Genes with a ratio smaller than 0.33, meaning export rate is at least twice the degradation rate, are considered to have no nuclear degradation (n = 5799), genes with ratio between 0.33 and 0.66 are considered low nuclear degradation (n = 2585) and genes with ratio above 0.66, meaning degradation rate is at least twice the export rate, are considered to have high nuclear degradation (n = 1478). (B) Results from gene set enrichment analysis, ranking genes according to share of nuclear degradation (see A). Hand-picked subset of significantly enriched gene ontology terms is shown. Color transparency shows negative logarithm to base 10 of Benjamini-Hochberg corrected false discovery rate (FDR). (C) Violin plots of share of nuclear degradation and nuclear retention, cytosolic, membrane and model-derived whole-cell half-lives for transcripts encoding transcription factors (n = 722), mitochondrial proteins (n = 826), ribosomal proteins (n = 96) and other (n = 8218). See Methods for gene group definitions. Center points of violin plots depict the median values.

**Supp. Fig. 5:**
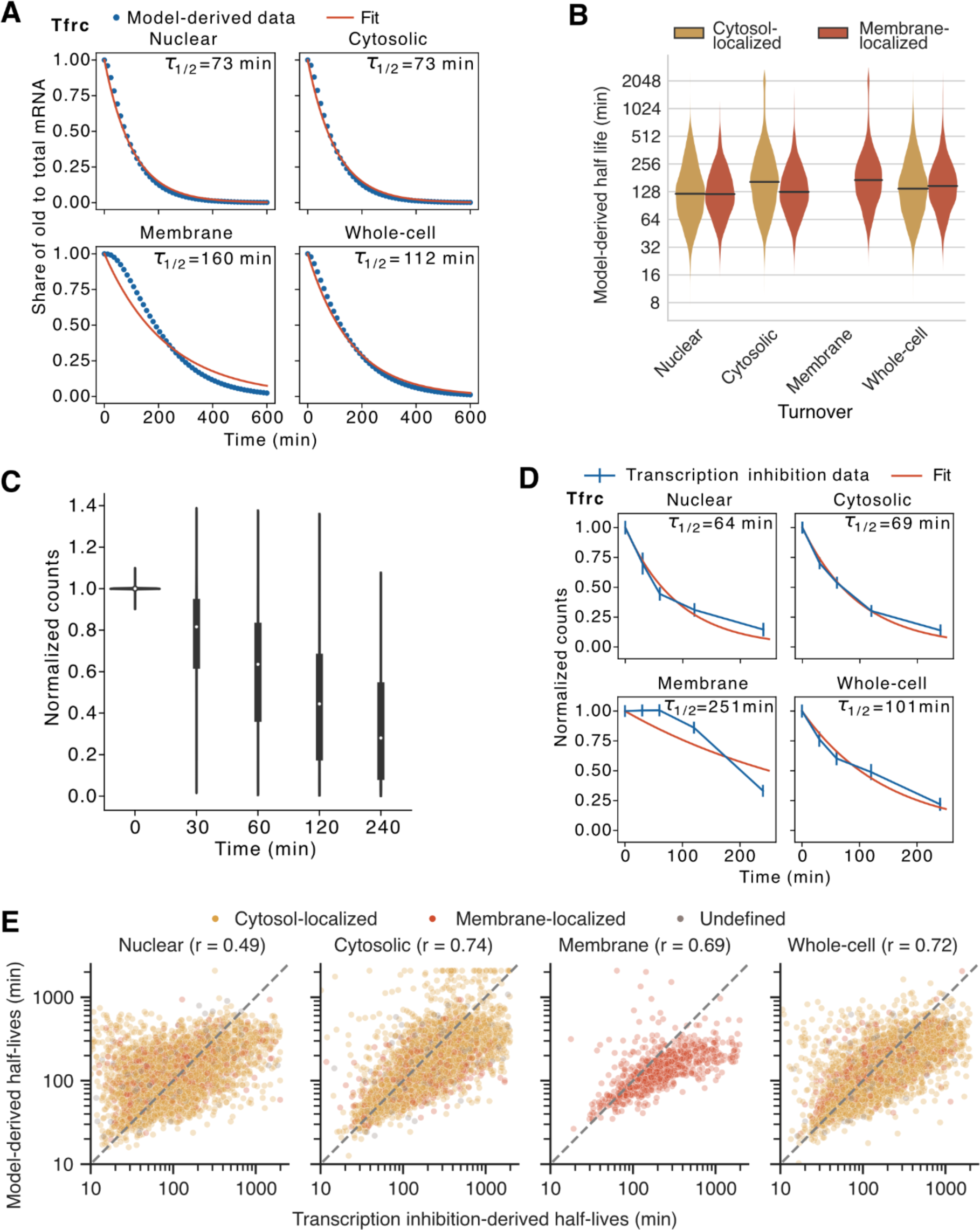
Estimating model-derived pulse-chase-like and transcription inhibition-derived mRNA half-lives. (A) Exemplary fit to determine model-derived, aggregated half-lives, shown for *Tfrc*. From the subcellular rates, model-derived pulse-chase trajectories are calculated for different compartments (blue dots). To this data, a simple exponential decay model is fit (red line). Best fit values are shown on top right in each parameter facet. (B) Violin plot showing half-lives of model-derived, aggregated parameters for cytosol-(yellow, n = 7818) and membrane-localized (red, n = 1635) transcripts. (C) Violin/box plot of normalized counts from transcription inhibition experiment using Flavopiriodol (all compartments). X-axis shows time exposed to flavopiridol. White dots depict median values. Lower and upper hinges of box plots correspond to the 25th and 75th percentiles, respectively. Upper and lower whiskers extend from the hinge to the largest or smallest value no further than the 1.5x interquartile range from the hinge, respectively. (D) Exemplary fit to determine transcription inhibition-derived half-lives, shown for *Tfrc*. To normalized counts (blue dots), a simple exponential decay model is fit (red line). Best fit values are shown on top right in each parameter facet. (E) Scatterplots of model-derived (x-axis) and transcription inhibition-derived (y-axis) half-lives of nuclear mature (n = 6018), cytosolic (n = 5966), membrane (n = 780) and whole-cell (n = 6340) parameters. Color indicates localization. Spearman rank correlation is shown in facet title. Dashed, gray line is the identity line.

